# Establishment of a Patient-Derived, Magnetic Levitation-Based, 3D Spheroid Granuloma Model for Human Tuberculosis

**DOI:** 10.1101/2021.04.01.438053

**Authors:** Leigh A. Kotze, Caroline G.G. Beltran, Dirk Lang, Andre G. Loxton, Susan Cooper, Maynard Meiring, Brigitte Glanzmann, Craig Kinnear, Gerhard Walzl, Nelita du Plessis

**Affiliations:** DST-NRF Centre of Excellence for Biomedical Tuberculosis Research; South African Medical Research Council Centre for Tuberculosis Research; Division of Molecular Biology and Human Genetics, Faculty of Medicine and Health Sciences, Stellenbosch University, Cape Town, South Africa.; Confocal and Light Microscopy Imaging Facility, University of Cape Town, Cape Town, South Africa.; South African Medical Research Council Genomics Centre.

## Abstract

Tuberculous granulomas that develop in response to *Mycobacterium tuberculosis* (*M.tb*) infection are highly dynamic entities shaped by the host immune response and disease kinetics. Within this microenvironment, immune cell recruitment, polarization and activation is driven not only by co-existing cell types and multi-cellular interactions, but also by *M.tb*-mediated changes involving metabolic heterogeneity, epigenetic reprogramming and rewiring of the transcriptional landscape of host cells. There is an increased appreciation of the *in vivo* complexity, versatility and heterogeneity of the cellular compartment that constitutes the tuberculosis (TB) granuloma, and the difficulty in translating findings from animal models to human disease. Here we describe a novel biomimetic *in vitro* 3-dimentional (3D) human lung granuloma model, resembling early “innate” and “adaptive” stages of the TB granuloma spectrum, and present results of histological architecture, host transcriptional characterization, mycobacteriological features, cytokine profiles and spatial distribution of key immune cells. A range of manipulations of immune cell populations in these granulomas will allow the study of host/pathogen pathways involved in the outcome of infection, as well as pharmacological interventions.

**IMPORTANCE:** Tuberculosis is a highly infectious disease, with granulomas as its hallmark. Granulomas play an important role in the control of M.tb infection and as such are crucial indicators for our understanding of host resistance to TB. Correlates of risk and protection to M.tb are still elusive, and the granuloma provides the perfect environment in which to study the immune response to infection and broaden our understanding thereof; however, human granulomas are difficult to obtain, and animal models are costly and do not always faithfully mimic human immunity. In fact, most TB research is conducted in vitro on immortalized or primary immune cells and cultured in 2D on flat, rigid plastic, which does not reflect in vivo characteristics. We have therefore conceived a 3D, human in vitro granuloma model which allows researchers to study features of granuloma-forming diseases, in an 3D structural environment resembling in vivo granuloma architecture and cellular orientation.

## INTRODUCTION

A hallmark of TB is the formation of granulomatous lesions in response to *M.tb*-infected phagocytes, such as alveolar macrophages (AM) within the pulmonary space, inducing a chronic inflammatory response. Additional macrophages are recruited to the site of infection to form the “core” structure of the granuloma, characterized as an innate response to infection. Recent literature shows that early *M.tb* infection occurs almost exclusively in airway-resident alveolar macrophages, whereafter *M.tb*-infected, but not uninfected, alveolar macrophages localize to the lung interstitium, preceding *M.tb* uptake by recruited monocyte-derived macrophages and neutrophils (Cohen et al., 2018). Other immune cell types such as interstitial macrophages, monocytes, dendritic cells, neutrophils, and lymphocytes (T- and B-cells) are recruited to the site of infection where they are collected and organised around the core to form mature granuloma structures to contribute to an adaptive immune response (Davis and Ramakrishnan, 2009; Lin et al., 2006; Wolf et al., 2007). Granulomas are thus the main site of the host-pathogen interaction, primarily aimed at preventing *M.tb* dissemination, but during immune dysregulation, can also function as niche for *M.tb* survival and persistence (Ehlers and Schaible, 2013).

During TB disease, the associated damage to host lung tissue, as evidenced by necrosis and cavitation, which is associated with bacterial persistence and the development of drug-resistant *M.tb*. Ultimately, TB patients harbour a range of granulomas, which may comprise a spectrum of solid non-necrotizing, necrotic and caseous granulomas, each with its own distinct microenvironment; these have been demonstrated to not be limited to active TB patients, but are also observed in animals/non-human primates with both latent and reactivation cases of TB (Flynn et al., 2011). The mere formation of granulomas is thus insufficient for infection control, and instead, must function with the appropriate combination of host control measures, for example the correct balance of pro- and anti-inflammatory response mediators. The outcome of the host response to infection may be beneficial in that some granulomas resolve completely (sterilizing cure), while others progress to caseation and rupture, allowing for the uncontrolled dissemination of *M.tb* into the surrounding tissue – both scenarios have been known to occur within the same individual (Barry et al., 2009). The heterogeneity of granulomas even within the same patient highlights the need to study each of these individual structures as a whole to assess host-pathogen interactions at an individual granuloma level. Progress in understanding *M.tb* pathogenesis at a patient granuloma-level has been poor due to the difficulty in obtaining fresh human TB granulomatous lung tissue. For this reason, patient-driven immune responses to *M.tb* have been mainly assessed in primary host immune cells, cultured *in vitro* as traditional cell cultures on plastic plates optimized for tissue culture. Such traditional cell cultures with uniform exposure to pathogen/immune mediators have served as important tools for studying host-*M.tb* interactions. However, these non-physiological conditions fail to recapitulate key elements of cells residing in the complex tissue microenvironment including cell concentration and types, cell motility, multicellular interactions, cell expansion, spatiotemporal kinetics and geometries (Duval et al., 2017; Nunes et al., 2019). Such discrepancies present a significant barrier to interpretation and translation of findings from basic science, vaccine immunogenicity-, drug efficacy- and prevention of infection studies, ultimately limiting the impact of TB research on human health.

While animal models have contributed significantly to our understanding of the mammalian immune system, numerous examples have shown that laboratory animal species do not faithfully or in full, mimic human immunity or TB disease (Fonseca et al., 2017; Singh and Gupta, 2018; Zhan et al., 2017). To overcome these obstacles, successful 3D *in vitro* models can contribute to the ethos of the 3Rs of animal research by replacing the use of animals in research with insentient material obtained from willing donors, thereby also promoting human relevance (Herrmann et al., 2019). Advanced *in vitro* models (derived from *in vivo* spatial information) that recapitulate the spatial interactions between recruited monocytes, T-cells and B-cells in the context of the human lung granuloma, are lacking, but are incredibly necessary for the future of TB research and the successful dissection of disease pathology (Elkington et al., 2019).

In this study, we utilised primary human alveolar macrophages retrieved from the site of disease and autologous adaptive immune cells isolated from the periphery, to successfully establish an *in vitro* 3D granuloma model of human TB using magnetic cell levitation and BCG infection. Magnetic cell levitation is a recently developed method used to generate tumour spheroids during which tumour cells are pre-loaded with magnetic nanospheres which electrostatically attach to cell membranes, to form multicellular spheroids suspended in culture via an external magnetic field. Nanospheres subsequently detach from the cells, allowing unsupported growth as the structure starts to mimic extracellular matrix (ECM) conditions (Souza et al., 2010). Here we describe an *in vitro* model which could be used to examine stages of granuloma formation, demonstrated here as early (innate) and late (adaptive) granuloma types. These complex multidimensional structures are investigated at morphological level and subsequently dissociated to assess single cell characteristics for comparison to traditional cell cultures prepared using identical cell types, ratios, and culturing conditions. We demonstrate, along with the method of construction, the possible applications of this model, including a wide range of immunological assays. This model has proven to be stable and reliable for the investigation of mycobacterial granulomas in a tissue culture setting, opening the door for expansion of this model into additional molecular biology avenues.

## METHODS

### PRIMARY PROCESSING OF THE 3D SPHEROID GRANULOMAS

#### Study Subjects

This proof-of-concept study enrolled three HIV-uninfected participants between the ages of 18 and 70 years who presented to the Pulmonology Division of Tygerberg Academic Hospital with clinical indications of TB. These largely included clinical and radiological abnormalities as determined by chest x-ray (CXR). Mycobacteria growth indicator tube culture (MGIT) and/or GeneXpert MTB/RIF was performed. Written informed consent was obtained from all participants, and a summary of their demographics is given (Table 1). The study design was approved by the Stellenbosch University Ethics Review Committee (IRB number N16/04/050).

**Table 1:**
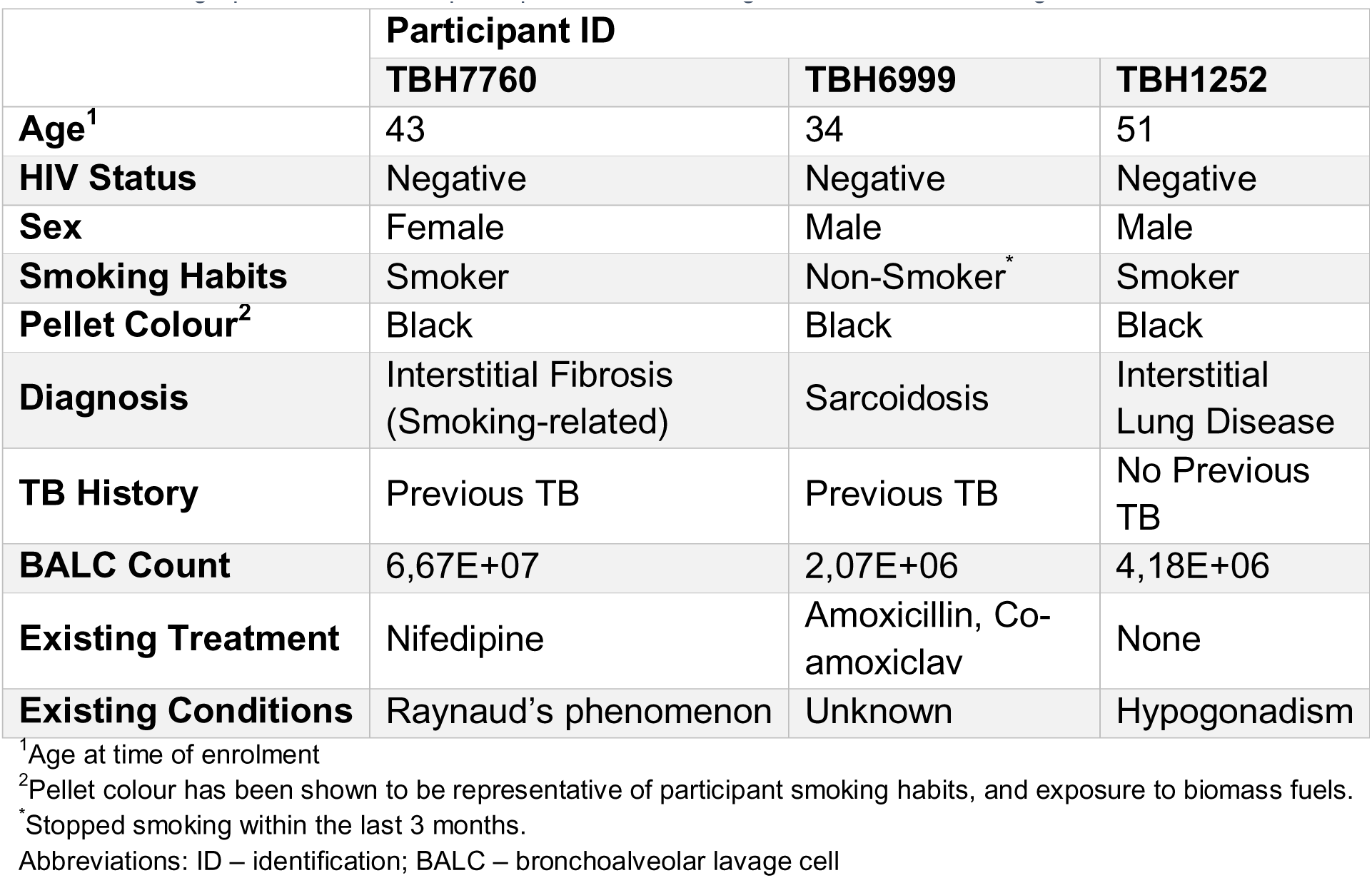
Demographic details of the participants used for the generation of *in vitro* 3D granulomas.

#### Sample Collection

A visual outline of the methods used for this study is shown in Figure 1.

**Figure 1:**
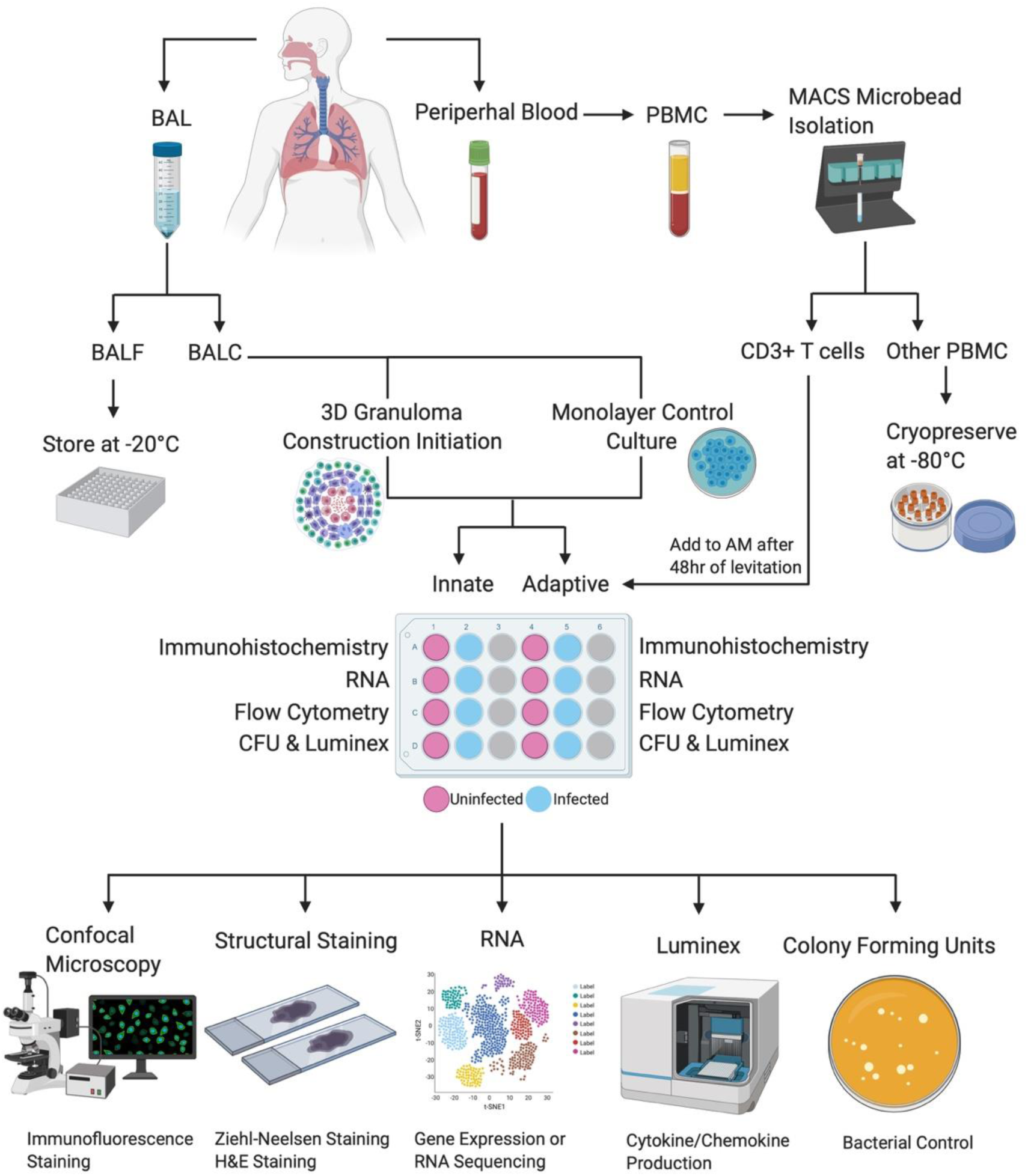
A basic representation of the workflow used to generate and analyse 3D *in vitro* human TB granulomas generated from a single participant. Briefly, BAL and peripheral blood are collected from the participant at the time of the bronchoscopy procedure, after which the BAL is processed to collect the cell pellet while the fluid is stored for other studies. The BAL cells (BALC) are then used to construct the alveolar macrophage core of the 3D structure and the traditional cell culture control (monolayer), in both uninfected and infected scenarios. Collected peripheral blood is processed for PBMC and then further processed to isolate autologous CD3^+^ T cells using the MACS MicroBead cell separation technique, with the CD3^-^ cellular fraction being stored for other studies. Autologous T cells are then added to the appropriate alveolar macrophage cores, those designated to become “adaptive” granuloma structures, after 48 hours of the core’s levitation or 48 hours of conventional culture in the case of the traditional cell culture control. Generated structures are then processed individually, in uninfected and infected pairs, for the respective downstream applications desired. These include embedding in tissue freezing media for subsequent cryosectioning and staining of the structures for immunofluorescence and confocal microscopy or staining for basic cellular structures like H&E staining or ZN staining for acid-fast bacterial detection (this is exclusively for the 3D structures and cannot be done for the traditional cell culture control cultures). Cells can also be stored for later RNA extractions and subsequent gene expression or RNA sequencing analyses. Supernatants can be stored for cytokine/chemokine production analyses using the Luminex immunoassay platform or similar platforms like ELISA, and cell lysates can be plated to determine CFU counts, thereby evaluating bacterial control.

Bronchoscopies were performed on the study participants by qualified clinicians and nursing staff according to international guidelines (Meyer et al. 2012a; Klech 1989) and bronchoalveolar lavage done in a segmental bronchus with radiologically suspicious lesions on CXR or computed tomography (CT). The lavage was performed by instilling sterile saline solution at 37°C up to a maximum volume of 300ml in aliquots of 60ml at a time, with aspiration between aliquots. Aspirated bronchoalveolar lavage (BAL) was collected in sterile

50ml polypropylene tubes (Falcon® 50ml Sterile Conical Centrifuge Tubes; Corning Inc., NY) and transported on ice to the laboratory. Immediately after bronchoscopy, peripheral blood samples were collected by venepuncture into two 9ml sodium heparinised (NaHep) vacutainers. Both BAL and peripheral blood samples were processed within 2 hours of collection and processed under BSL3 and BSL2 conditions, respectively, owing to the high infectious risk of BAL samples.

#### Cell Isolations

BAL cells (BALC) were isolated by centrifugation of BAL fluid (BALF) for 7 minutes at 300 x g (4°C) following sterile filtration through a 70μm cell strainer (Falcon® 70μm Cell Strainer; Corning Inc., NY) and successive wash steps with 1 X phosphate buffered saline (PBS). All cells were counted, and the viability determined using the Trypan Blue (0.4%; Thermo Fisher Scientific Inc., Waltham, MA) exclusion method (Strober 1997). BALC were cultured overnight to isolate adherent cells (the majority of which are alveolar macrophages (AM)) from non-adherent cells (lymphocytes) should the fraction contain lymphocytes exceeding 30% of total cells, and then cryopreserved in cryomedia consisting of 10% dimethyl sulfoxide (DMSO; Sigma-Aldrich Co., St. Louis, MO) and 90% fetal bovine serum (FBS; HyClone, GE Healthcare Life Sciences, Illinois, USA).

Peripheral blood mononuclear cells (PBMC) were isolated from peripheral blood by centrifugation using Ficoll density gradient medium (Histopaque-1077; Sigma Chemical Co., St. Louis, MO). Cells were counted and the viability determined using the Trypan Blue exclusion method. PBMC were then used to isolate autologous CD3^+^ T cells using the MACS→ (magnetically activated cell sorting) MicroBead Isolation technique (Miltenyi Biotec, Cologne, Germany) according to the manufacturer’s instructions. Both the CD3^+^ T cells and the CD3^-^ cell fractions were cryopreserved in cryomedia (described above).

#### Thawing of Cryopreserved Cells

Cryopreserved cells were thawed in a waterbath (37°C) and transferred to 10ml of warmed complete RPMI-1640 media (supplemented with 1% L-glutamine (Sigma-Aldrich, St Louis, Missouri, USA) and 10% FBS). Cells were centrifuged at 300 x g for 10 minutes and washed twice. All cells were counted, and the viability determined using the Trypan Blue exclusion method.

#### AnaeroPack Experiment

BALC were seeded in a 24-well low adherence plate at a concentration of 2×10^6^ cells per 500µl complete RPMI. A 2ml screw cap tube containing 2×10^6^ BALC in 500µl complete RPMI was cultured separately and acted as the untreated control. 100µM Cobalt Chloride (CoCl_2_; Sigma-Aldrich; catalogue number:C8661-25G) was added to each well to stabilize HIF-1α, before placing the plate in a lock-seal box. Working quickly, the AnaeroPack (Mitsubishi Gas Chemical Co. INC) was removed from the protective foil and placed within the lock-seal box next to the culture plate, and the box quickly sealed shut. The AnaeroPack System is an easy anaerobic atmosphere cultivation method that has no need for either water or catalyst. AnaeroPack creates an environment of <0.1% oxygen, and >15% CO_2_. Cells were incubated for 24 hours (37°C, 5% CO_2_), after which 125µl 16% paraformaldehyde (PFA; ThermoFisher Scientific, catalogue number: 28908) was immediately added to the well and untreated control tube (to make a final concentration of 4% PFA) after opening the lock-seal box. The box was re-sealed allowing for the cells to fix at room temperature for 20 minutes. Cells were transferred out of the plate to a 15ml Falcon tube and centrifuged at 400 x g for 10 minutes. The supernatant was removed, and the cells resuspended in 1ml 1XPBS. Cytospins of 2×10^5^ BALC were then created for both the treated and untreated control cells on poly-L-lysine-coated microscope slides (Sigma-Aldrich, Catalogue number: P0425-72EA), to be used for immunofluorescent staining.

#### BCG Infection

Cultures of *Mycobacterium bovis* Bacille Calmette-Guerin (BCG) were grown in Difco Middlebrook 7H9 (BD Pharmingen, San Diago, USA) supplemented with 0.2% glycerol (Sigma-Aldrich), 0.05% Tween-80 (Sigma-Aldrich), and 10% Middlebrook Oleic Acid Albumin Dextrose Catalase (OADC) enrichment (BD Pharmingen, San Diago, USA). Aliquots of BCG cultures at an OD_600_ of 0.8 were stored at −80°C in RPMI-1640 media supplemented with 10% glycerol (Sigma-Aldrich). The number of viable bacteria was assessed by thawing a frozen aliquot and plating serial dilutions onto Middlebrook 7H11 (BD Pharmingen, San Diego, USA) agar plates. The plates were incubated at 37°C (5% CO_2_) for 21 days and colonies counted manually thereafter to determine the number of viable bacteria.

AM were thawed and rested at 37°C (5% CO_2_) for 18 hours in complete RPMI-1640 supplemented with 1% L-glutamine (Sigma-Aldrich, St Louis, Missouri, USA), 10% FBS and an antimycotic antibiotic containing 10,000U/ml penicillin, 10,000μg/ml streptomycin and 25μg/ml amphotericin B (Fungizone) (PSF; Lonza, Walkersville, MD, USA). The following day, AM were washed in complete RPMI lacking PSF, centrifuged at 300 x g for 10 minutes and then cultured at a density of 4×10^5^ cells/well in a 24-well low adherence culture plate (Greiner Bio-One, North Carolina, USA). Both “Uninfected” and “Infected” wells were seeded for comparison between infection states. “Infected” wells were infected with BCG (Pasteur strain, MOI 1) and incubated for 4 hours. Extracellular bacteria were removed by incubating cells (37°C, 5% CO_2_) with complete RPMI supplemented with PSF for 1 hour, followed by successive washes with complete RPMI lacking PSF.

#### 3D Granuloma Construction

Magnetic cell levitation and -bioprinting are recently developed methods used to generate 3D tumor spheroids (Tseng et al., 2015, 2013). We have implemented this system to create incipient, innate-style and mature, classic (adaptive)-style 3D lung granuloma types resembling TB granulomas, which will henceforth be referred to as innate and adaptive granulomas, respectively. It should be noted that all incubation periods mentioned in this methods section are carried out in an incubator (37°C, 5% CO_2_), and that paired BALC and PBMC samples were used to build each spheroid.

In brief, BCG uninfected and infected AM were treated overnight with biocompatible NanoShuttle^TM^ (made from gold, iron oxide and poly-l-lysine; n3D Biosciences Inc., Greiner Bio-One) at a concentration of 100µL per 1×10^6^ cells in low-adherence plates at a maximum volume of 300µL per well (Haisler et al., 2013; Souza et al., 2010). The following day, the NanoShuttle^TM^-labelled AM were levitated using the magnetic levitating drive (n3D Biosciences Inc., Greiner Bio-One) for 48 hours to cluster and assemble the granuloma core (Figure 2a). This granuloma core consisting of only AM is considered the innate granuloma. Concurrently, autologous CD3^+^ T cells were thawed, counted, and rested overnight in complete RPMI in a separate 24-well culture plate. Adaptive granulomas were created 48 hours post-construction of the granuloma core, through addition of these autologous T cells (6×10^5^/well) at a ratio of 40:60 (AM:T cells). We previously conducted a titration of cell ratios and selected a macrophage to lymphocyte ratio that macroscopically resembles a TB lung granuloma considered to harbour a median bacterial burden (Wong et al., 2018). This was achieved by carefully removing the levitating drive from the culture plate (Figure 2b) and replacing it with the magnetic printing drive (Figure 2c). Chemokine production by the AM core allows for T cell migration to the granuloma structure, forming an outer lymphocytic cuff, closely resembling the complex *in vivo*, organized mature classic human TB granulomas. The adaptive granuloma was then incubated for 24 hours, after which the printing drive was removed, and each granuloma dedicated to various downstream processes. For the innate granulomas, no autologous T cells were added but the core was incubated for the same period as the adaptive granulomas (Figure 2d). Nanospheres subsequently detach from the cells, allowing unsupported growth as the structure starts to mimic extracellular matrix (ECM) conditions. This model thus allows for the addition, enrichment, or depletion of specific innate or adaptive immune cells or immune mediators to interrogate their potential contribution to protective or nonprotective granulomatous responses.

**Figure 2:**
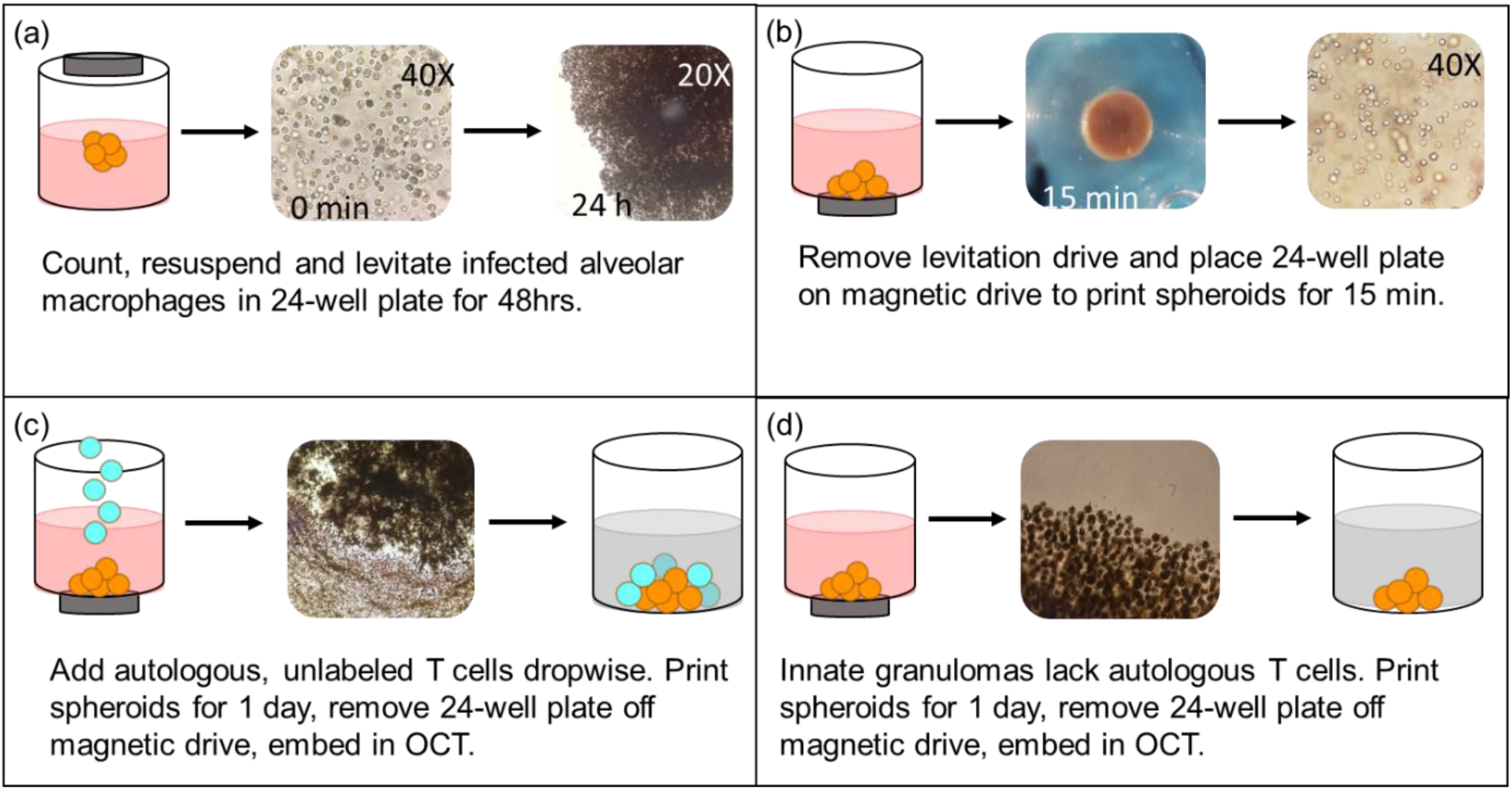
3D Innate and Adaptive Granuloma construction using the n3D Biosciences Inc. magnetic levitation and printing drives. (a) The development of the granuloma core is accomplished through the levitation of NanoShuttle^TM^-labelled alveolar macrophages for 48 hours. (b) The magnetic levitation drive is removed after 48 hours and immediately replaced by the magnetic printing drive below the 24-well culture plate which ensures that the 3D structure remains intact. (c) Autologous CD3+ T cells, not labelled with NanoShuttle^TM^, are then carefully added to the alveolar macrophage core, and allowed to migrate via chemotactic gradients to the core to create the mature, adaptive granuloma. (d) Innate granulomas do not have the autologous CD3+ T cells added to the core but are rather left with the printing drive secured below the culture plate for the remainder of the experiment.

#### Traditional Cell Culture Control Construction

As a control culture condition, traditional cell cultures were constructed in tandem with the 3D granuloma construction in a separate 24-well low adherence culture plate for each participant. The exact same ratios of AM to T cells were used for these control cultures, the only difference being that **no** NanoShuttle^TM^ was added to the AM, and subsequently no levitation- or printing drives were added to the culture plate to replicate cellular distributions under normal *in vitro* culture conditions. Both “Uninfected” and “Infected” control cultures were set up, for both innate and adaptive granuloma types.

### DOWNSTREAM PROCESSING OF THE 3D SPHEROID GRANULOMAS

#### 3D Granuloma Embedding and Cryosectioning

A pair of Uninfected and Infected granulomas had their supernatant carefully removed and stored in separate 0.2mL Eppendorf tubes at −80°C for later cytokine response interrogation using Luminex immunoassays. The structures were then washed twice with 300µL complete RPMI, making sure to not disrupt the structures, and subsequently fixed with 300µL 4% paraformaldehyde (PFA; ThermoFisher Scientific, Catalogue number: 28908) for 30 minutes in the dark. Fixed structures were then embedded in tissue freezing media, OCT (Tissue-Tek; USA), and stored at −80°C for later cryosectioning. Cryosections were made using a −20°C Cryostat (Leica Biosystems), with sections being cut 7µM thick and mounted onto poly-L-lysine-coated microscope slides (Sigma-Aldrich, Catalogue number: P0425-72EA). The cryosections were numbered in order of cutting to allow for the “position” within the structure to be inferred and picked for specific staining. Mounted sections were stored at −20°C until immunohistochemistry staining could be performed. Sections were reserved for immunofluorescent staining (sections from the centre of the 3D granuloma structure), H&E staining (a section from near the centre) and ZN staining (a section from near the centre). The processing described for “Immunofluorescent Staining”, “Immunofluorescent Microscopy”, “H&E Staining”, “Ziehl-Neelsen Staining” and “Light Microscopy” was not done for the traditional cell culture conditions.

#### Staining and Microscopy

##### 1. Immunofluorescent Staining

All staining was done in a humidified chamber. Fixed granuloma sections (stored at −20°C) were rehydrated in 1 X PBS for 10 minutes followed by two washes of 1 X PBS for 5 minutes. Sections were blocked with 1% Bovine Serum Albumin (BSA; Sigma-Aldrich, Catalogue number: A7030) in PBS containing 0.1M glycine for 1 hour at room temperature after which the sections were incubated with corresponding primary antibodies diluted in 1% BSA in PBS overnight at 4°C in the dark. Antibody concentrations were as follows: alveolar macrophage membranes were stained using CD206 PE-CF594 (Mouse Anti-Human CD206, BD Biosciences; catalogue number: 564063) at a concentration of 4µL in a 50μL final staining volume per section; and the universal T cell surface marker CD3 V450 (Mouse Anti-Human CD3, BD Biosciences; catalogue number: 560365) was used at a concentration of 5μL in a 50µL final staining volume per section. To assess necrosis, the expression of the necrotic marker, HMGB1, was assessed using Mouse Anti-Human HMGB1 (BioLegend, catalogue number: 651408) at a concentration of 20µg/mL. HMGB1 stained sections were counterstained with the nuclear stain, Hoechst (Life Technologies; catalogue number: H3570), at a concentration of 1µg/mL. Following overnight incubation, sections were washed three times in 1 X PBS for 5 minutes and the sections allowed to dry slightly before mounting with DAKO mounting medium (Agilent Technologies) and allowing to airdry overnight in the dark at room temperature. Slides were stored at 4°C in the dark until imaging. Unstained and single stained controls were also prepared for each experiment to assess background and signal specificity in each channel.

Immunofluorescent staining for cytospin slides (see “**AnaeroPack Experiment**” Methods) were performed as described above, with antibody concentrations as follows: BALC were stained with Hoechst (Life Technologies; catalogue number: H3570) at a concentration of 1 μg/mL; the hypoxia marker HIF-1α AF488 (Mouse Anti-Human, BioLegend; catalogue number: 359708) was used at a concentration of 2.5μg/mL final staining volume per section; and the necrosis marker HMGB1 AF647 (Mouse Anti-Human, BioLegend; catalogue number: 651408) was used at a concentration of 20μg/mL final staining volume per section.

##### 2. Immunofluorescent Microscopy

Images were obtained using a Carl Zeiss LSM 880 Airyscan with Fast Airyscan Module confocal microscope (Plan-Apochromat x63/1.40 oil DIC UV-VIS-IR M27 objective lens), and the images were acquired using the ZEN software (Carl Zeiss). Acquisition settings for imaging were identically set for all sections. Red channel: excitation wavelength (561nm), emission wavelength (659nm), detection wavelength (585-733nm), pinhole (1.42 AU), frame scan mode, detector gain (640); blue channel: excitation wavelength (405nm), emission wavelength (432nm), detection wavelength (410-455nm), pinhole (2.17 AU), frame scan mode, detector gain (724). 18×18 tile scans were acquired and stitched together.

##### 3. H&E Staining

One section from both Uninfected and Infected granuloma structures, not reserved for immunofluorescent staining and control slides, was stained using the Lillie Mayer H&E method (Lillie, 1965) to visualise the overall compartmentalisation of the various cell types added to the 3D structure without immunofluorescent stains. Briefly, the designated sections were removed from storage and allowed to reach room temperature while placed in a slanted position and rehydrated in dH_2_O. Sections were then stained with alum haematoxylin for 4 minutes, rinsed with tap water and differentiated with 0.3% acid alcohol for 2 seconds. Sections were rinsed again with tap water, then rinsed in Scott’s tap water substitute, and rinsed again with tap water. Finally, sections were stained with Eosin for 2 minutes, dehydrated, cleared and coverslips mounted using 1 drop of VectaMount^TM^ (Vector Laboratories, California, USA).

##### 4. Ziehl-Neelsen (ZN) Staining

One section from both Uninfected and Infected granuloma structures, not reserved for immunofluorescent staining and control slides, was stained using the ZN method to identify the localisation of acid-fast bacteria within the 3D structure. Briefly, the designated sections were removed from storage and allowed to reach room temperature while placed in a slanted position. Sections were then flooded with carbol fuchsin and heated gently with a flame until a vapour is emitted. Sections were immersed for 5 minutes, rinsed with dH_2_O, flooded with acid-alcohol, and allowed to stand for 2 minutes. Following this, sections were rinsed with dH_2_O, counterstained with methylene blue, allowed to stand for 2 minutes and finally rinsed and allowed to dry. Dry sections were mounted with coverslips using 1 drop of VectaMount^TM^.

##### 5. Light Microscopy

H&E-stained sections and ZN-stained sections were visualised using the ZEISS Axio Observer microscope, fitted with an Axiocam MRc 195 microscope camera, and the images were acquired using the ZENlite imaging software (blue edition, version 1.1.1.0).

#### Flow cytometry

An Uninfected and Infected granuloma pair was mechanically dissociated (by gentle pipetting) into single cells and transferred to respective sterile 15mL Falcon tube. Cells were then centrifuged at 300 x g for 10 minutes and resuspended in 5mL complete RPMI and washed. Washed cells were then resuspended in 300µL complete RPMI and transferred to a sterile, 24-well culture plate (untreated), after which cells were incubated for 48 hours to allow for adherence of the AM. Following incubation, the non-adherent fraction was separated from the adherent cells and the wells washed twice. These cells were immediately used for the flow cytometric investigation of the phenotypic and functional characteristics of the non-adherent cellular fraction, namely T cells. The same procedure was followed for an Uninfected and Infected traditional cell culture control pair. The adherent fraction was not analysed by flow cytometry, due to the high levels of autofluorescence. To demonstrate this, 1×10^6^ total BALC and PBMC were transferred to two separate 5ml Falcon tubes and centrifuged at 250 x g for 5 minutes. Both fractions were kept unstained and acquired on the BD FACS^TM^ Canto II, with gates being set using a pre-defined FMO from a stained PBMC sample with the channels CD3 PE-Cy7, CD14 Pacific Blue, and anti-HLA-DR APC.

Non-adherent fractions were analysed by flow cytometry using the BD FACS^TM^ Canto II, and were stained to define their basic phenotypic profiles. Fractions were stained using anti-CD3 PE-Cy7 (BD Biosciences; catalogue number: 563423), anti-CD14 Pacific Blue (BD Biosciences; catalogue number: 558121), anti-CD16 PerCP-Cy5.5 (BD Biosciences; catalogue number: 565421), and anti-CD56 BV510 (BD Biosciences; catalogue number: 563041). Staining was performed for 30 minutes in the dark at room temperature, in a 50μl staining volume. Titrations, compensation, and fluorescence-minus-one (FMO) for each antibody was performed prior to analysis at the BD Stellenbosch University Flow Cytometry Unit on Tygerberg Campus. Data was analysed using the third party FlowJo Software v10.0.8. and the 3D granuloma structure data compared to that obtained for the traditional cell culture control.

#### RNA Sequencing

##### 1. RNA Extractions

An Uninfected and Infected granuloma pair was mechanically dissociated (by gentle pipetting) into single cells and transferred to a sterile 2mL screw-cap tube. Cells were then centrifuged at 300 x g for 10 minutes and vigorously resuspended in 350µL RLT Buffer (Qiagen, Germany) by vortexing for 30 seconds. Cells were stored at −80°C for batched RNA extractions to perform gene expression or RNASeq analysis. On the day of batched RNA extractions, samples were removed from storage and allowed to thaw at room temperature. RNA was extracted using the Qiagen RNeasy® Mini Kit according to the manufacturer’s instructions. Following isolation, 1µL of each sample was used to check RNA integrity using a Nanodrop spectrophotometer (Thermo Fisher Scientific, Massachusetts, USA), with all samples having a A260/A280 ratio of above 1.8. The remaining RNA was stored until RNA Sequencing could be performed. The same procedure was followed for an Uninfected and Infected traditional cell culture control pair.

##### 2. Library Preparation

Total RNA was extracted from eleven samples; however, two samples did not meet the total RNA (200ng) requirements for further processing. Total RNA was subjected to DNase treatment and magnetic bead-based mRNA enrichment using the Dynabeads™ mRNA Purification kit (Invitrogen™, Thermo Fisher Scientific, Waltham, MA, USA), according to the protocol described in the *MGIEasy RNA Library Prep Set User Manual* prior to proceeding with library construction. Library preparation was performed with the entire component of mRNA for each sample using the *MGIEasy RNA Library Prep Kit* (MGI, Shenzen, China), according to the manufacturer’s instructions.

##### 3. Sequencing

Massively parallel sequencing was performed at the South African Medical Research Council (SAMRC)/Beijing Genomics Institute (BGI) Genomics Centre using DNA nanoball-based technology on the MGISEQ-2000 using the appropriate reagents supplied in the MGISeq-2000RS High-Throughput Sequencing Kit. A paired-end sequencing strategy was employed with a read length of 100bp (PE100). Sample libraries were loaded onto MGISEQ-2000 FCL flow cells with the MGILD-200 automatic loader, and 18 FASTQ files generated (9 forward- and 9 reverse read files).

##### 4. Analysis

Raw FASTQ files were assessed using FastQC (version 0.11.5). Reads with quality scores less than 20 and length below 30 bp were all trimmed. The resulting high-quality sequences were subsequently used in MultiQC, a module contained in Python (version 3.6.3), to aggregate and summarise the results from multiple FastQC reports into a single HTML report. Raw FASTQ files were then imported, annotated (human GRCh38.p13 dataset from https://www.ncbi.nlm.nih.gov/assembly/GCF_000001405.39/), filtered (Counts per Million (CPM) cut-off method), normalised, and analysed using statistical software based in R (version 3.6.3). Differentially regulated transcripts were functionally annotated to gain an overview of biological pathway regulation. Briefly, GO terms enrichment analysis was conducted on the ensemble gene IDs using the Database for Annotation, Visualization and Integrated Discovery (DAVID) v 6.8 (Huang et al., 2009a, 2009b), while the REVIGO resource was used to summarise and visualize the most enriched GO terms (Supek et al., 2011).

#### Luminex® Immunoassay

As mentioned previously in the methods section, a pair of Uninfected and Infected granulomas had their supernatant removed and stored in separate 0.2mL Eppendorf tubes at −80°C for later cytokine response interrogation using the Luminex immunoassay platform (Luminex, Bio Rad Laboratories, Hercules, CA, USA). Supernatants were collected and stored from structures throughout the culture period, beginning from 1-day to 4-days post infection, at the beginning of each day’s processing. The same procedure was followed for an Uninfected and Infected traditional cell culture control pair. A one-plex kit was used to measure the production of interleukin (IL)-22 (R&D Systems®, LXSAHM-01), and a four-plex was used to measure the production of IL-2, IL-10, interferon (IFN)-γ, and tumor necrosis factor (TNF)α (R&D Systems®, LXSAHM-04). Briefly, samples were removed from storage and allowed to reach room temperature one hour before beginning the assay. Samples were then vortexed and prepared for the assay according to the manufacturer’s instructions. Samples were not diluted as recommended by the manufacturer but run neat due to the small number of cells used in culture and the restrictive lower limit of detection observed for Luminex immunoassays. Samples were run on the Luminex® MAGPIX system. The beads from each sample were acquired individually and analysed using the Bio-Plex Manager^TM^ Software version 6.1 according to recommended settings. Instrument settings were adjusted to ensure 50 bead events per region, with sample size set to 50μl for both kits.

#### CFU Determination

The supernatant from the remaining Uninfected and Infected granuloma pair was carefully removed and the cells mechanically dissociated (by gentle pipetting) into single cells after adding 300µL complete RPMI. Cells were then transferred to a sterile 15mL Falcon tube and centrifuged at 300 x g for 10 minutes and resuspended in 200µL dH_2_O to lyse the cells and vortexed. Serial dilutions were prepared using this cell lysate using PBS-Tween80. The neat cell lysate and serial dilutions were then plated out in duplicate on Middlebrook 7H11 agar plates (BD Biosciences) for manual CFU determination after 21 days. The same procedure was followed for an Uninfected and Infected traditional cell culture control pair.

#### Statistical analysis

Statistical analyses were performed using GraphPadPrism version 8 (GraphPad Software, San Diego, CA). A p-value of less than 0.05 was considered significant. Tests for normality could not be performed due to sample sizes being too small in this pilot study; data was, therefore, treated as nonparametric throughout. Where two groups were compared the Mann-Whitney t-test was used. The Kruskal-Wallis test was performed using Dunn’s post-test to correct for multiple comparisons where three or more groups were compared.

## RESULTS

The results of this proof-of-concept study demonstrate that the *in vitro* 3D human granuloma model is capable of being successfully interrogated using multiple molecular biology platforms.

### Macroscopic and Microscopic Differences in Granuloma Structural Formation exist due to cell composition and mycobacterial infection

NanoShuttle^TM^-labelled alveolar macrophages were levitated for 48 hours at the beginning of the model construction, and granuloma cores were observed as hanging cultures with spheroids suspended just underneath the surface of the culture media. After the first 24 hours of levitation, the magnetic levitation drive was removed briefly to investigate the structural integrity of the granulomas without magnetic influence. The assembly of the AM core could already be visualised at the 24-hour time point with BCG infected cores appearing less “stable” than uninfected cores (Figure 3a), with this apparent instability improving by the 48-hour culture time point (Figure 3b). At the end of the culture period, slight macroscopic variations in the morphology of uninfected (Figure 3c) and BCG infected (Figure 3d) innate granulomas could be seen, as well as between the uninfected (Figure 3e) and BCG infected (Figure 3f) adaptive granulomas. A “cuff” of unlabelled autologous CD3^+^ T cells were clearly seen to be surrounding the darker, NanoShuttle^TM^-labelled AM core (Figure 3e and 3f), compared to the innate core lacking the “cuff” due to the lack of autologous CD3^+^ T cells (Figure 3c and 3d). While these visual differences are important for defining structural phenotypes during the early stages of this model’s development, information regarding details like the infiltration of autologous CD3^+^ T cells into the AM core could not be inferred without immunofluorescent microscopic assessment.

**Figure 3:**
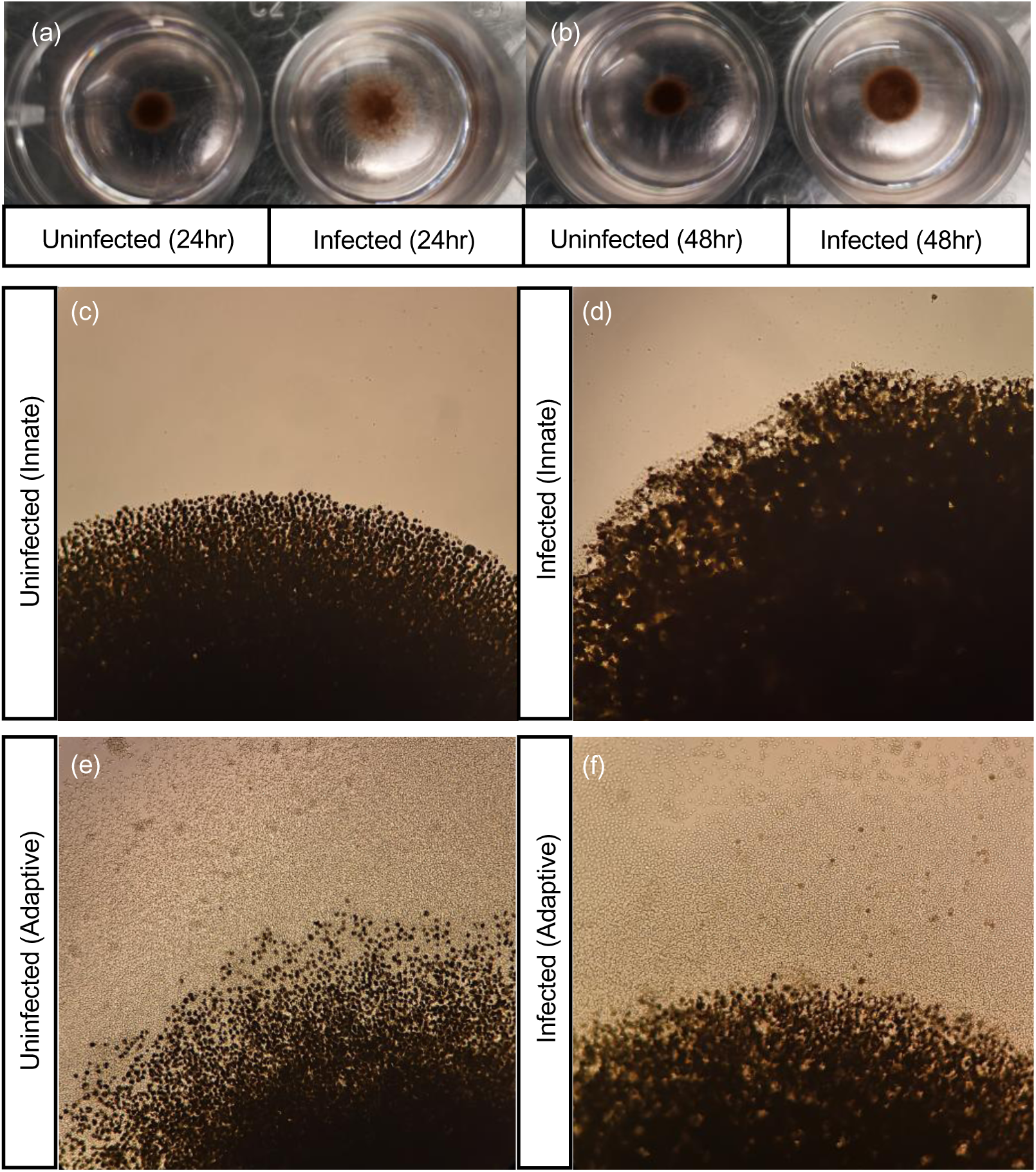
Visual differences in cellular organisation between uninfected and BCG infected granuloma structures can be observed after (a) 24- and (b) 48-hours of magnetic levitation, with BCG infected structures displaying less robust structural integrity (the magnetic levitation drive was removed briefly for these images to be taken). Differences in cellular organisation could also be visualised using light microscopy, with (c) uninfected and (d) BCG infected innate granuloma structures displaying a clear lack of a lymphocytic cuff at the end of the culture period. Both the (e) uninfected and (f) BCG infected adaptive granuloma structures displayed the presence of a lymphocytic cuff (unlabelled, clear cells) surrounding the NanoShuttle^TM^-labelled AM core (darker cells) at the end of the culture period during which time both magnetic levitation and magnetic bioprinting was used. Images were taken using an inverted light microscope (20x magnification).

Granuloma structures were further investigated at cellular level to interrogate the composition in a more detailed manner. For the uninfected granulomas, both H&E (Figure 4c) and ZN (Figure 4d) stains demonstrated that no acid-fast bacilli are present within the AM core (Figure 4d). ZN staining of an infected granuloma structure demonstrated acid-fast bacilli both near the cuff of the structure (Figure 4i) and in the core (Figure 4j). For both infected and uninfected granulomas, the adaptive granuloma structures demonstrated a “cuff” of autologous lymphocytes, based on known morphology, as seen using both H&E (Figure 4g) and ZN (Figure 4h) staining. It is interesting to note that the core of the adaptive granuloma structures was almost exclusively made up of macrophages (Figure 4c), suggesting minimal infiltration by autologous CD3^+^ T cells from the “cuff” at this infection time point. Inferences could suggest that the architecture of granulomas with non-infiltrating lymphocytes encapsulating the AM core, represent a structural component of mycobacterial infection control/failure (Gil et al., 2010; Lenaerts et al., 2015; Peyron et al., 2008); alternatively, this could also be a methodological limitation whereby the CD3^+^ T cells have simply not had sufficient time to migrate through the core.

**Figure 4:**
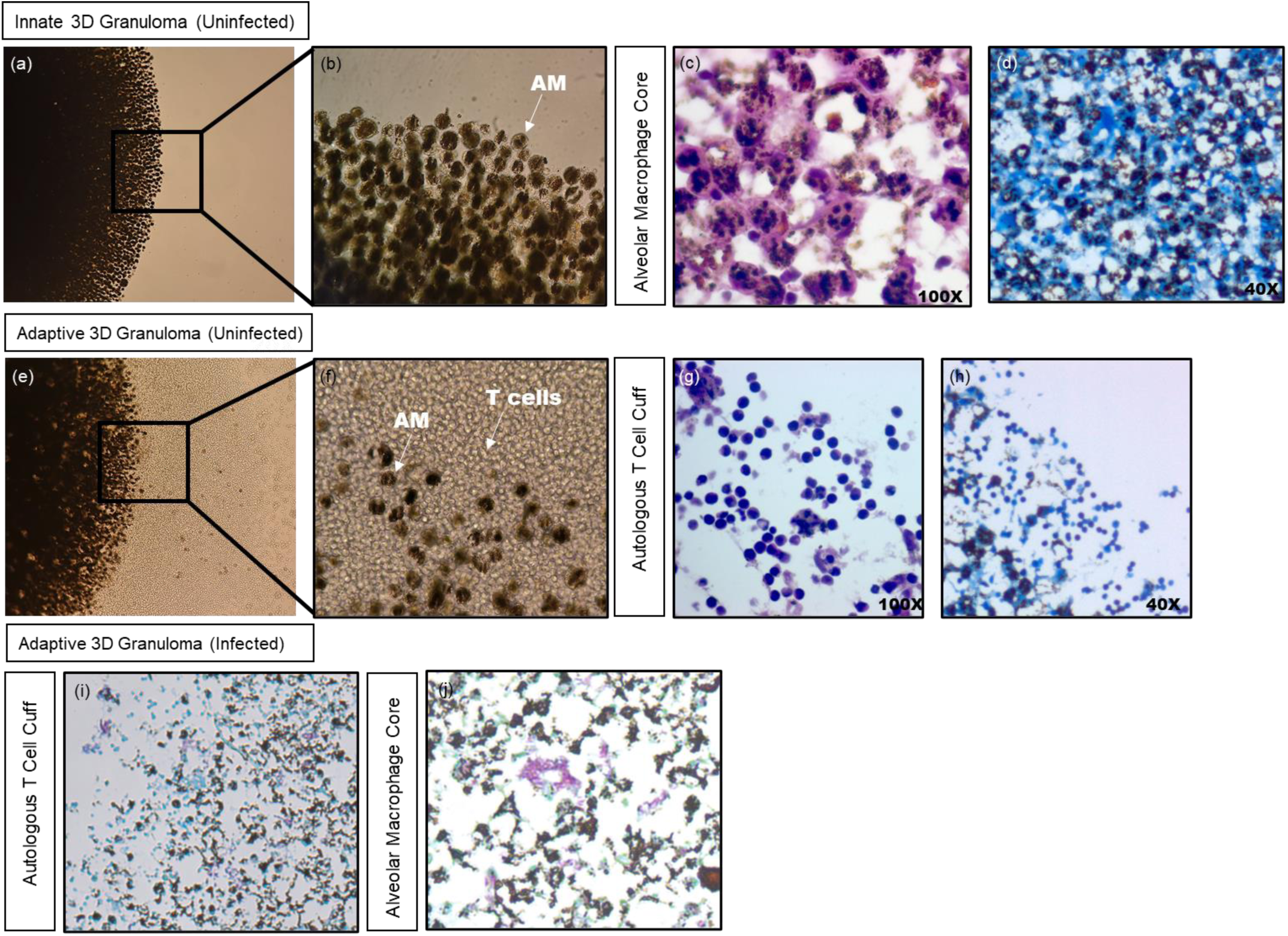
Demonstration of the various staining method outcomes performed using sections from an uninfected innate ((a)-(d)) and adaptive ((e)-h)) 3D granuloma structure. Light microscopy of the (a) innate and (e) adaptive 3D granuloma structures clearly demonstrate the (b) alveolar macrophage core, with (f) CD3+ autologous T cells forming a cuff around the core in the adaptive granuloma structure. These observations are corroborated using the H&E staining method, once again demonstrating (c) the alveolar macrophage core and (g) CD3+ autologous T cell cuff. ZN staining of the sections demonstrated the uninfected nature of the structures, both in (d) the core and (h) the cuff, while ZN staining of infected sections demonstrated acid-fast bacilli both near (i) the cuff and (j) in the core of the structure.

### Magnetic levitation combined with bioprinting, successfully produces a morphologically and physiologically relevant 3D TB lung granuloma

Confocal microscopy (Figure 5a) demonstrated that both the outer edges (Figure 5b) and core (Figure 5c) of the innate granuloma is made up exclusively of AM (CD206 PE-CF594 (red)). A tile scan (Figure 6a) of the adaptive granuloma mid-section revealed a “cuff” of autologous CD3^+^ T cells (Figure 6b), and an exclusively AM-dominant core (Figure 6c), again demonstrating little-to-no autologous CD3^+^ T cells infiltrating the core. These findings are synonymous with the inferences made about human granuloma structures from non-human primate (NHP) models of TB (Flynn et al., 2015).

**Figure 5:**
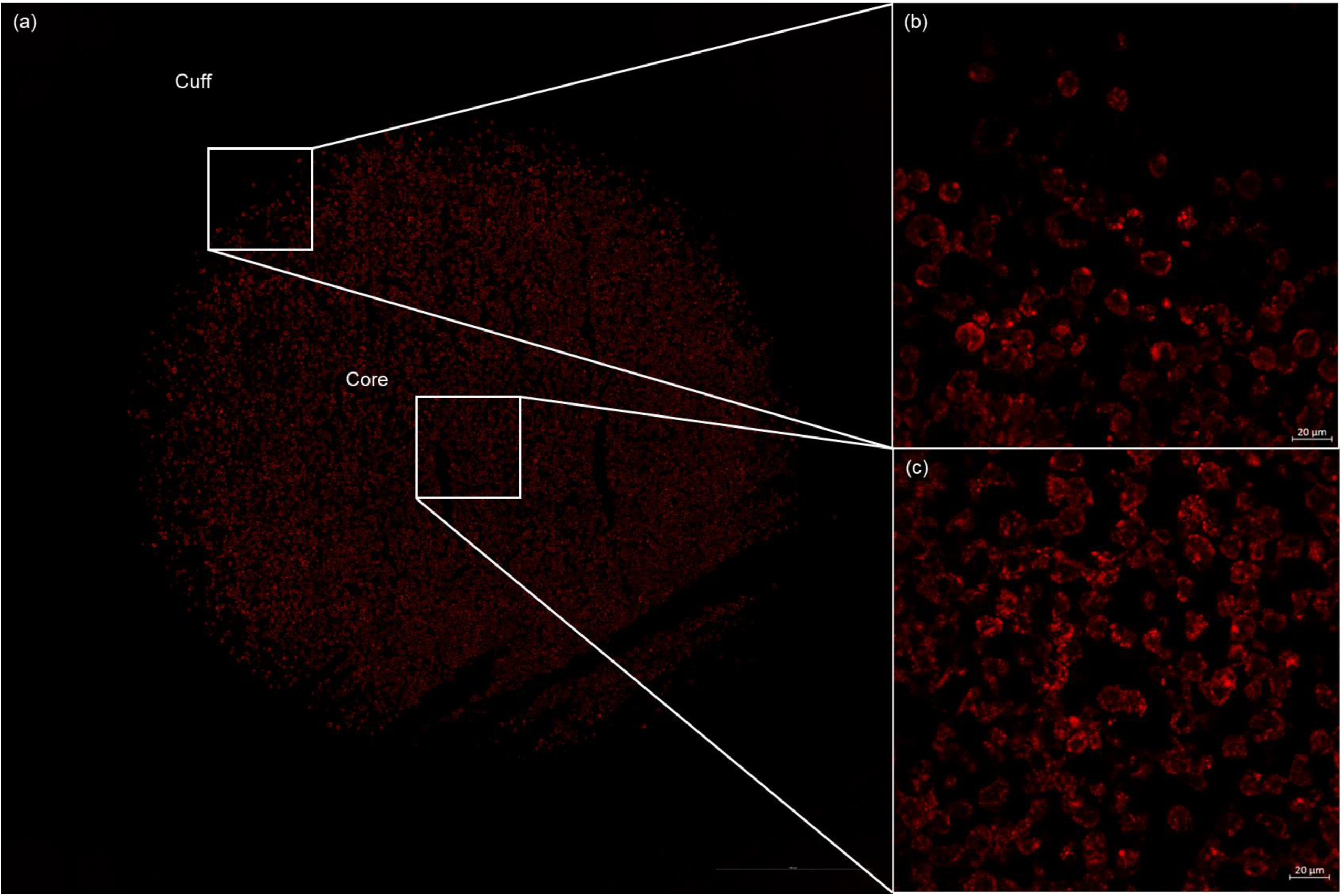
Tile scan (14×14) of a 3D uninfected innate granuloma structure depicting (a) the entire structure, (b) the “cuff” devoid of autologous CD3^+^ T cells, and (c) the exclusively AM-dominant core. AM were stained with CD206 PE-CF594 (red). Nuclei were left unstained.

**Figure 6:**
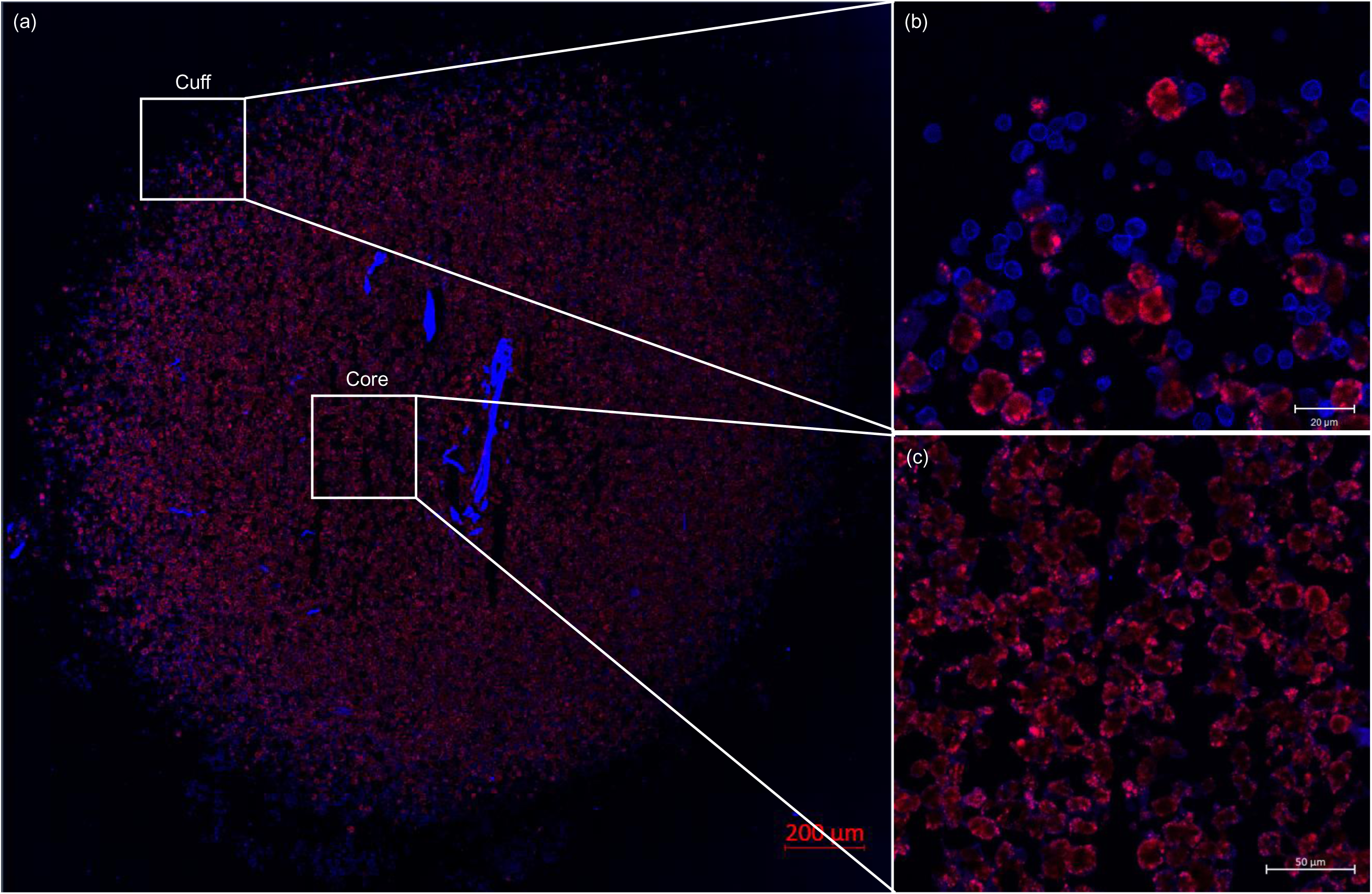
Tile scan (18×18) of a 3D uninfected adaptive granuloma structure depicting (a) the entire structure, (b) the autologous CD3^+^ T cell-dominant “cuff”, and (c) the exclusively AM-dominant core. AM were stained with CD206 PE-CF594 (red) and autologous CD3+ T cells were stained with CD3 V450 (blue). Nuclei were left unstained.

In addition to our 3D spheroid granuloma structures being morphologically relevant, it remains important to use this model to recapitulate physiologically relevant events of *in vivo* granulomas. It is well known that TB granulomas undergo structural and localised changes during development and maturation. These changes often include the caseation of the core through the process of necrosis which contributes to morbidity by causing tissue damage. The caseous granuloma *in vivo* is notably hypoxic, driving the accumulation of hypoxia inducible factor 1-α (HIF-1α), and considered the hallmark of failed *M.tb* containment, implicated in transmission (Belton et al., 2016; Canetti, 1955; Pagán and Ramakrishnan, 2014). HIF-1α, which is ubiquitously expressed in the cytoplasm under normoxic conditions, is used in this study as a hypoxia target (Hirschhaeuser et al., 2010), while necrotic cells are known to release the high mobility group box 1 (HMGB1) protein from the nucleus into the cell cytoplasm (Parasa et al., 2014). We compared the expression of HMGB1 and HIF-1α in our 3D spheroid granuloma model to traditional control cultures. Traditional cell culture controls created using BAL cells in an AnaeroPack experiment displayed basal expression patterns for both targets, synonymous with cells being non-nectrotic and normoxic (Figure 7a). Following experimental manipulation of traditional cultures to induce a hypoxic environment (Figure 7b), an upregulation in cytoplasmic expression of HIF-1α, although no nuclear translocation (indicative of a hypoxic cell) was observed (Figure 7c); however, the cytoplasmic translocation (indicative of a necrotic cell) of HMGB1 from the nucleus to the cytoplasm of BAL cells was achieved in traditional culture after experimental induction (Figure 7d). In contrast, we investigated both targets within our 3D spheroid granuloma model (Figure 7e), which did not require any additional experimental manipulation whatsoever such as was necessary for the traditional cell culture controls. Here we demonstrated that cytoplasmic translocation of HMGB1 successfully occurs in BAL macrophages when located at the core of the 3D spheroid granuloma (Figure 7f) and remains in the nucleus of T-cells at the cuff (Figure 7g), displaying a brightly stained rim. We were unsuccessful in capturing any HIF-1α immunofluorescent signals within the 3D spheroid granuloma model and we propose that this is as a result of HIF-1α being highly ubiquitinated upon re-exposure to oxygen, making it difficult to target without knowing the point at which hypoxia is induced during *M.tb* infection (Salceda and Caro, 1997). Our data confirms that the core of the 3D spheroid granuloma contains AM, which are both necrotic and non-necrotic, and a cuff of non-necrotic T-cells surrounds this core, recapitulating both morphological and physiological characteristics of TB granulomas. While traditional cell cultures require experimental manipulations for the induction of hypoxic and necrotic responses seen *in vivo*, our 3D human spheroid granuloma can recapitulate the *in vivo* granuloma microenvironment without external manipulation, by mimicking the spatial organization and cellular interactions within the 3D conformation, which may allow for the appropriate investigation of these forms of cell death.

**Figure 7:**
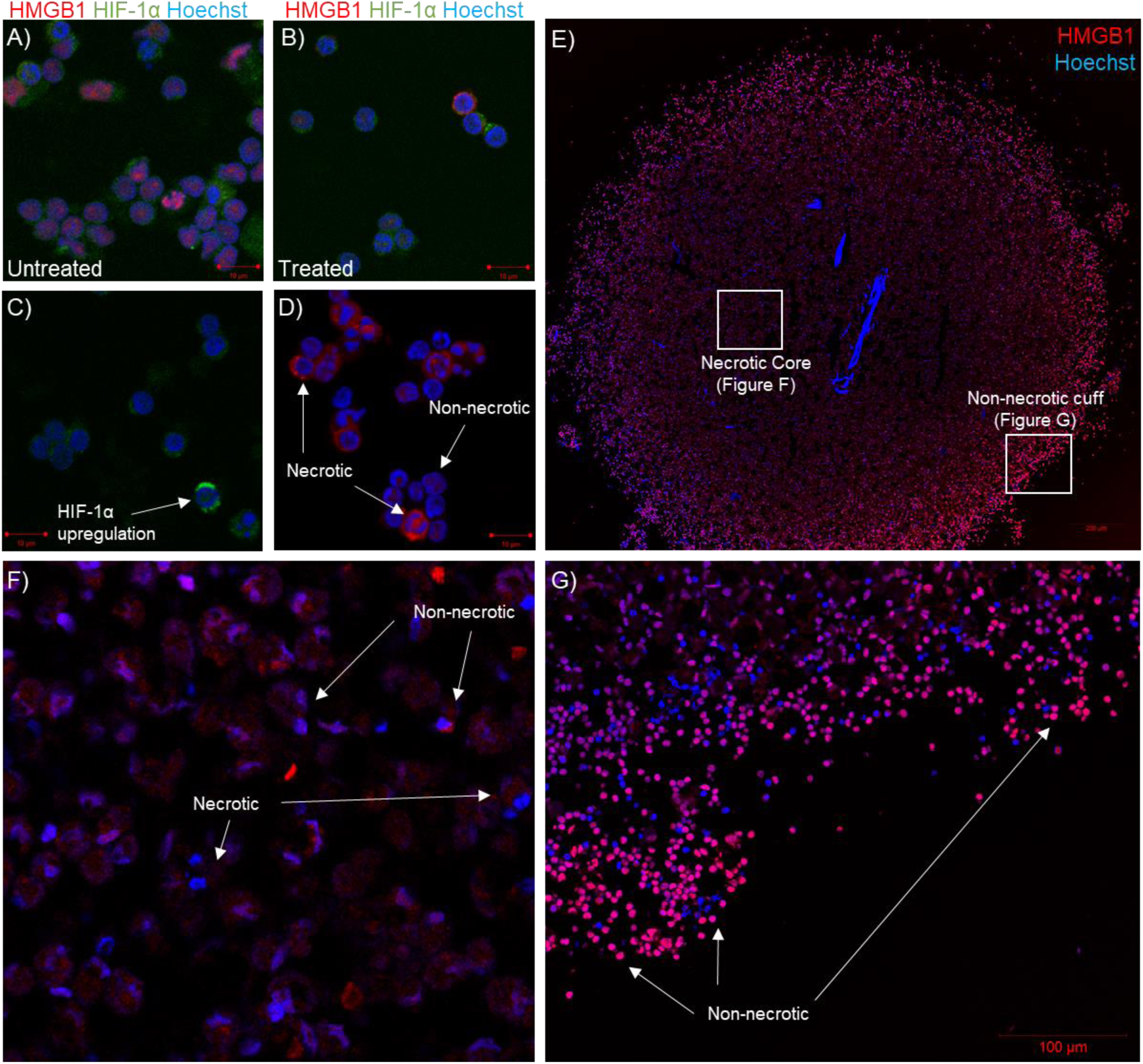
Confocal microscopy imaging of BAL cells **A)** under basal culture conditions (untreated) and **B)** under experimentally induced hypoxic conditions (treated), stained with HMGB1 (red), HIF-1α (green) and Hoechst (nuclei). Single stains of **C)** the HIF-1α and **D)** the HMGB1 markers under chemically induced hypoxic conditions demonstrated unsuccessful capturing of HIF-1α nuclear translocation during hypoxia, but successful capturing of cytoplasmic translocation of HMGB1 proteins, indicative of necrosis. Confocal microscopy imaging of **E)** the entire 3D human TB granuloma section stained with HMGB1 (red) demonstrated the establishment of an oxygen gradient resulting in **F)** an AM core of both necrotic and non-necrotic cells, and **G)** a cuff of non-necrotic cells.

### Single cell immune phenotyping of post-culture granuloma cells reveals maintained viability and distinct immune phenotypes

The viability and cell number of mechanically disrupted 3D granuloma structures was assessed via flow cytometry and compared to traditional cell culture controls. Innate granuloma structures could not be assessed in this manner owing to the highly auto-fluorescent nature of AM, making flow cytometry near-impossible on these cells (Figure 8). Particulate matter found within the AM of smokers or persons exposed to biomass fuels appear to be responsible for the autofluorescence, as is demonstrated by signals observed in the red channel during lambda scanning on the confocal microscope (Figure 9) (Young et al., 2019). Importantly, we could demonstrate that it was the AM and not the CD3^+^ T cells which were autofluorescent. Therefore, only adaptive granuloma structures were mechanically dissociated after the full culture period of six days as these could then be cultured for adherence to separate the AM from the T cells. Following prolonged culture, we have shown that the AM core becomes necrotic, and thus expect the cells to dye more easily with Trypan blue compared to traditional cell culture controls. Using the Trypan blue exclusion method, cell viability (Figure 10a) and cell number (Figure 10b) could be assessed and compared for each structure. No statistically significant differences were observed between the four groups for either cell viability or cell counts based on the small sample size.

**Figure 8:**
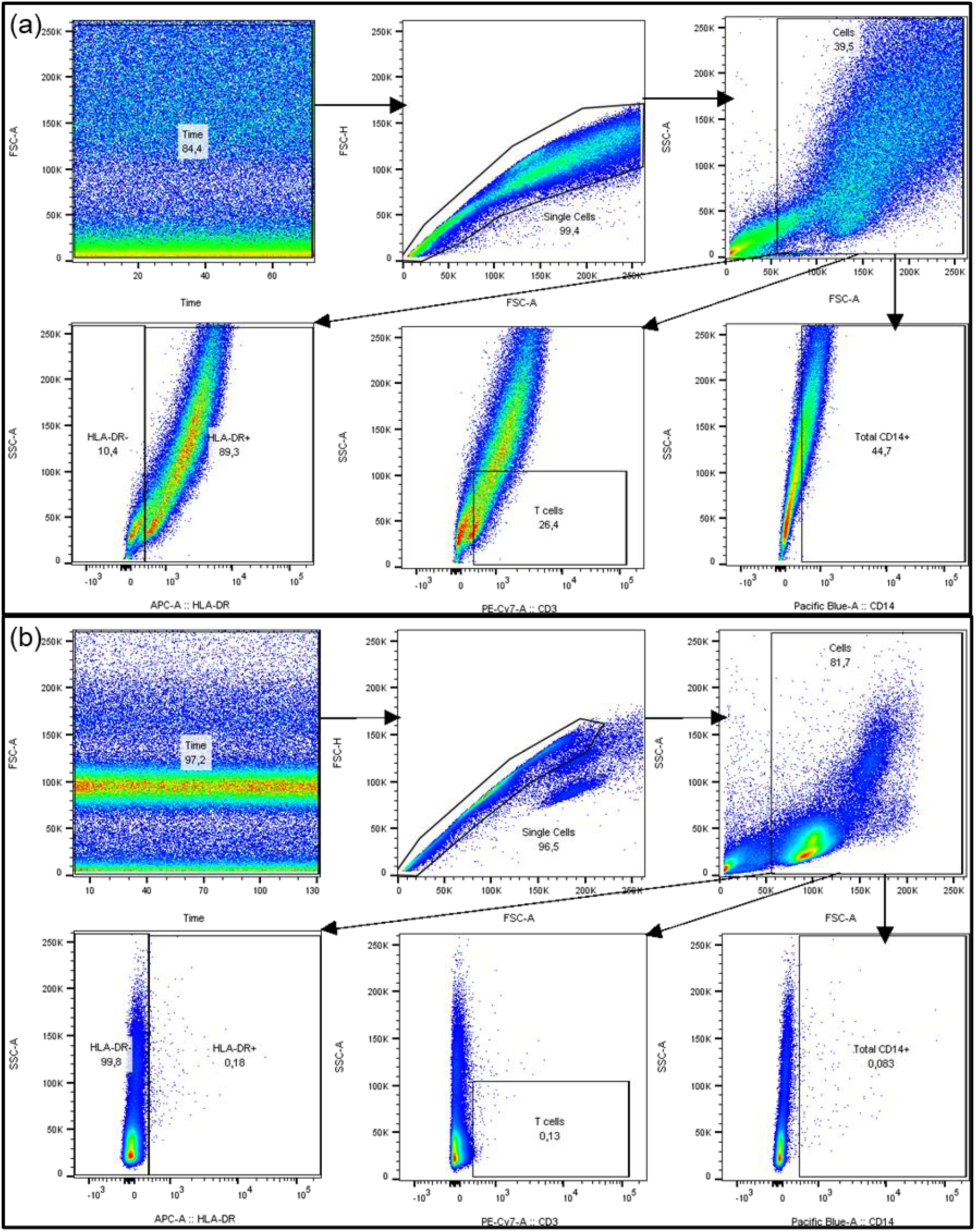
(a) Unstained total BALC run on the BD FACS™Canto II demonstrate positive autofluorescent signals compared to (b) unstained PBMC. Gates were set using FMO quality control checks set using a PBMC sample.

**Figure 9:**
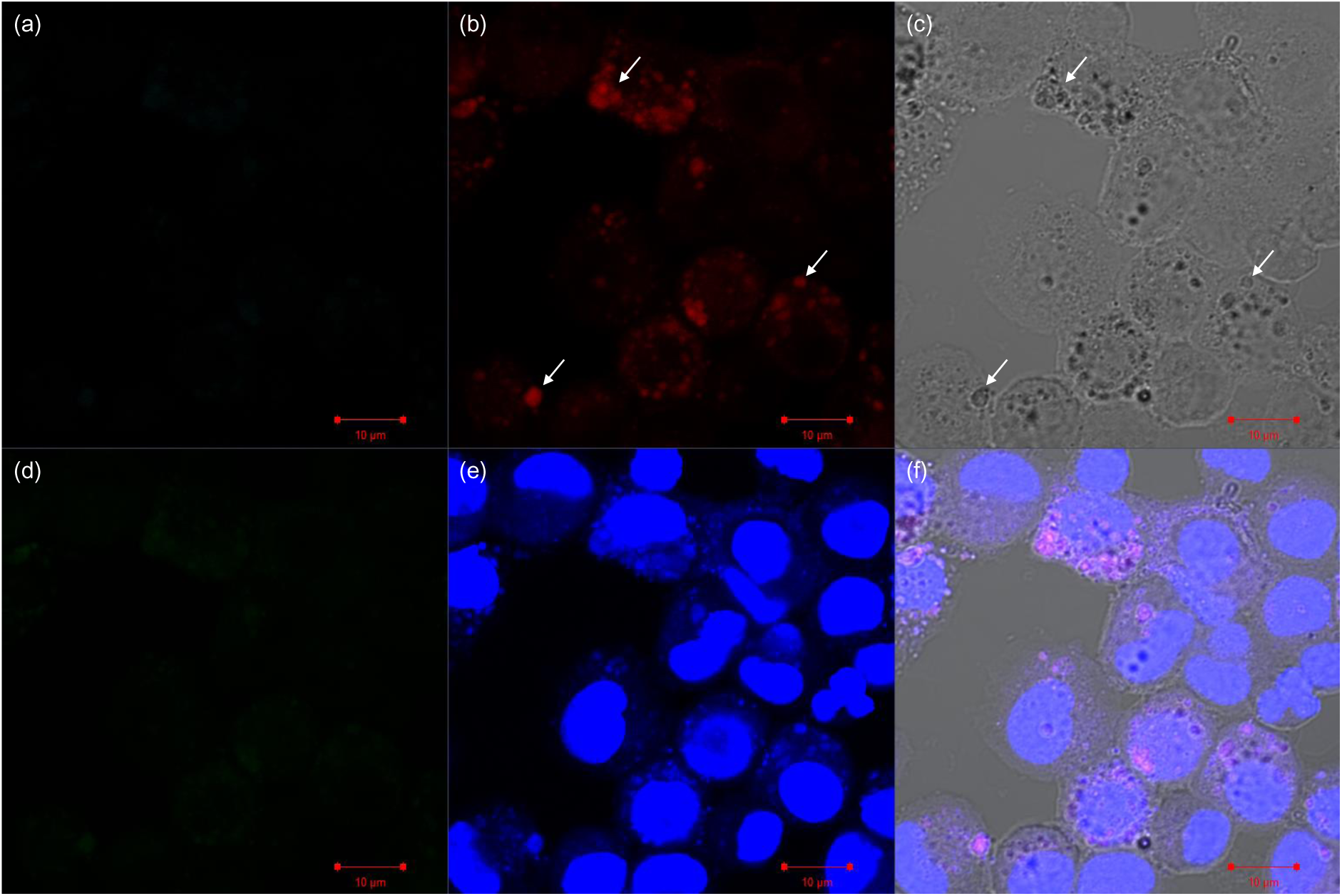
Unstained alveolar macrophages stained with Hoechst (nuclear stain) alone for slide orientation and focus. Auto-fluorescent signal was detected in the red (b), green (d) and blue (e) channels, with the red signal originating from the particulate matter-containing vesicles identifiable under brightfield (c) and the overlay (f). White arrows indicate particulate matter-containing vesicles.

**Figure 10:**
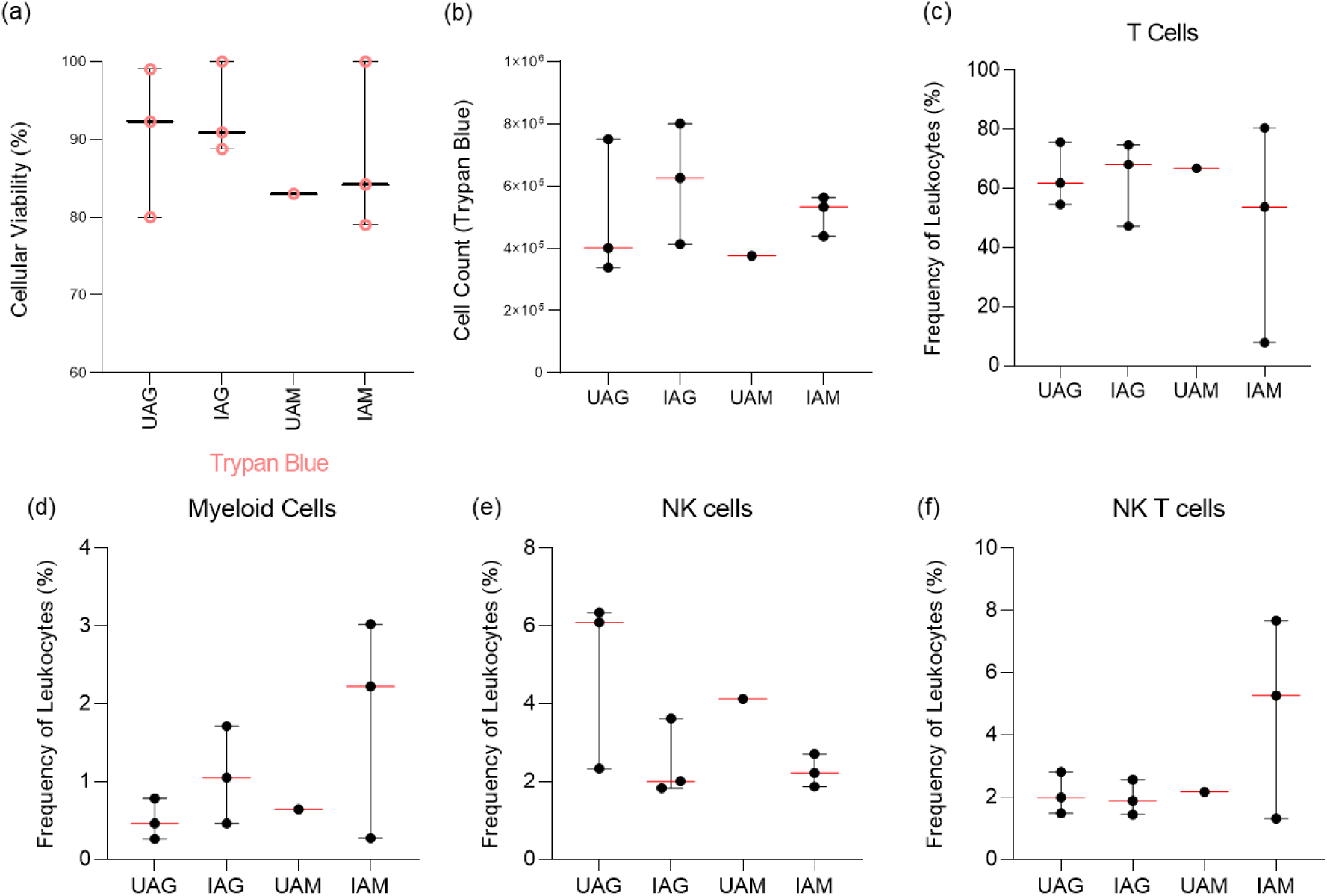
From each granuloma structure from each participant, the viability of mechanically dissociated granuloma structure single cells could be determined using the (a) Trypan Blue exclusion method (pink, circular datapoints). (b) Viable cells could be counted using the Trypan Blue exclusion method, and using flow cytometry, the frequency of various immune cell phenotypes could be assessed after the 48-hour adherence period to remove AM, including (c) CD3^+^ T cells, (d) CD3^-^CD14^+^ Myeloid Cells, (e) CD3^-^CD16^+^ Natural killer (NK) cells, and (f) CD3^+^CD16^+^ NK T cells. Cell counts and viability data are representative of cellular integrity after the final 48-hour incubation for adherence for both uninfected- (n = 3) and infected- (n = 3) granuloma structures, as well as uninfected- (n = 1) and infected- (n = 3) traditional cell cultures (monolayer). Each point represents a single data point; error bars are representative of the median and range. Abbreviations: UAG – Uninfected adaptive granuloma; IAG – Infected adaptive granuloma; UAM – Uninfected adaptive monolayer; IAM – Infected adaptive traditional.

We then investigated the ability to conduct single cell immune phenotyping on deconstructed granuloma single cells following six days of culture. We also wanted to know how the 3D conformation and resultant molecular signals, would impact cell frequencies, when compared to traditional cell culture cultures. From the same mechanically dissociated adaptive granuloma structures and corresponding traditional cell culture controls as used for the assessment of viability, phenotyping by way of flow cytometry (FACS^TM^ Canto II) could be successfully performed on the non-adherent cell fraction. Autologous CD3^+^ T cells (Figure 10c), CD3^-^CD14^+^ Myeloid cells (Figure 10d), CD3^-^CD16^+^ NK cells (Figure 10e), and CD3^+^CD16^+^ NKT cells (Figure 10f) could be identified from the non-adherent cell fraction, with CD3^+^ T cells proving to be the dominant cell subset. While none proved statistically significant (Table 2), there were visible differences in the phenotypic profiles of the non-adherent cell fractions of the uninfected- and infected-adaptive granuloma structures, as well as the uninfected- and infected-traditional cell culture controls (referred to as a “monolayer” within the figure). While the proof-of-concept nature of this study does not allow for strong biological inferences made based on the results observed for the granuloma structures and traditional cell cultures, our study does show promise for future investigations where we plan to significantly increase the number of granuloma structures assessed.

**Table 2:** Mann-Whitney t-test results for comparisons between infection groups for each cellular phenotype, as depicted above in Figure 3.

| **Comparison\*** | **T Cells**<br>**(p-value)** | **Myeloid Cells**<br>**(p-value)** | **NK Cells**<br>**(p-value)** | **NKT Cells**<br>**(p-value)** | **NK Regulatory**<br>**Cells (p-value)** |
| --- | --- | --- | --- | --- | --- |
| **Uninfected**<br>**Adaptive**<br>**Granuloma vs**<br>**Infected Adaptive**<br>**Granuloma** | **>0,9999** | **0,3** | **0,2** | **0,7** | **0,7** |
| **Infected Adaptive**<br>**Granuloma vs**<br>**Infected Adaptive**<br>**Traditional cell**<br>**culture** | **>0,9999** | **0,7** | **>0,9999** | **0,7** | **0,4** |

\*Comparisons could not be made between the Uninfected Adaptive Granuloma and Uninfected Adaptive Monolayer, and Uninfected Adaptive Monolayer and Infected Adaptive Monolayer groups due to too few datapoints being available for the Uninfected Adaptive Monolayer group.

### 3D granulomas secrete pro-inflammatory cytokines into the extracellular environment, signalling the recruitment of other immune cells

Traditional cell cultures are known for their capabilities of producing cytokines and chemokines during short term cultures, as measured frequently by Luminex immunoassays. Supernatants from 3D granuloma structures and traditional cell culture controls were harvested throughout the culture period, and at the end of the six-day culture. The most interesting differences to investigate would be the production of the selected cytokines from both AM and autologous CD3^+^ T cells, comparing these between 3D granuloma structures and traditional cell culture controls. We performed a Luminex immunoassay on the harvested supernatants and measure the concentrations of IL-10 (Figure 11a), TNF-α (Figure 11b), IFN-γ (Figure 11c), IL-2 (Figure 11d) and IL-22 (Figure 11e). Tests for normality could not be performed due to the small sample size, but QQ plots of the data showed the data could follow a Gaussian distribution if more datapoints were available. Data was, therefore, treated as nonparametric data, with the Kruskal-Wallis test being performed using Dunn’s post-test to correct for multiple comparisons. All cytokines assessed could be successfully measured from the harvested supernatants, with the cytokine IFN-γ showing high production in T cells prior to their addition to the granuloma structure, as would be expected from a normal culture. While none of the cytokines showed statistically significant differences between the groups, this was expected due to the low sample size, but was performed regardless to demonstrate the viability of the generated 3D granuloma structures over a long culture period and interact in a manner reminiscent of the *in vivo* human TB granuloma. Of particular interest was the observation that the innate granulomas (both uninfected (UIG) and BCG infected (IIG)) did not produce high levels of cytokines like TNF-α, IFN-γ and IL-2 associated with T cell production and release, further supporting the previous statement. Whilst not significantly different, the differences between the 3D granuloma structures and the traditional cell controls could prove to be higher in the 3D granuloma structures should we increase the sample size, as a few differences were already apparent, for example IFN-γ.

**Figure 11:**
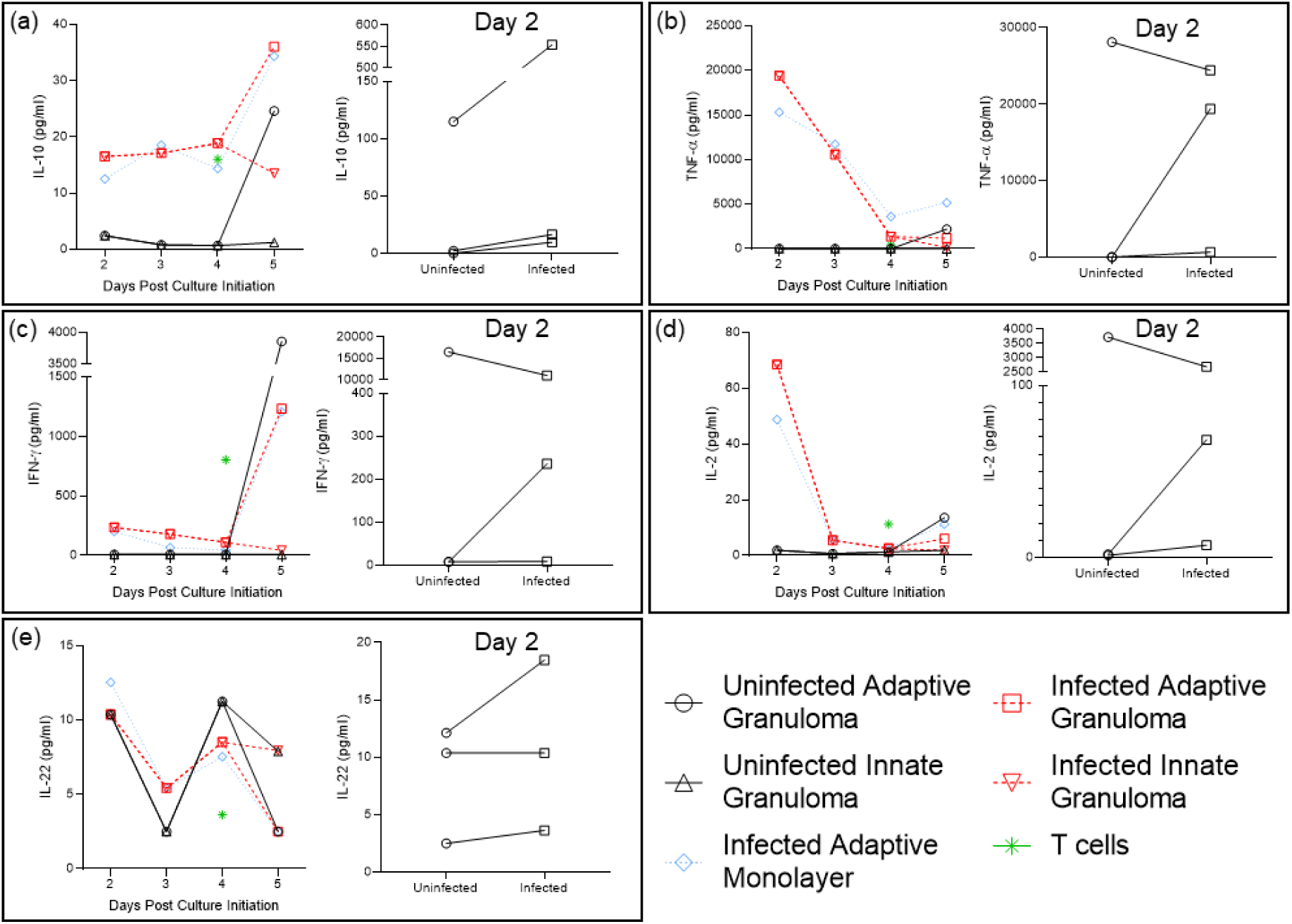
Concentration (pg/ml) of the investigated cytokines released into the supernatant of the 3D granuloma structure and traditional cell culture control (monolayer) extracellular environments, as measured by Luminex analysis. The cytokines measured included (a) IL-10, (b) TNF-α, (c) IFN-γ, (d) IL-2, and (e) IL-22. Cytokine production was compared between initial production by uninfected and BCG infected AM 2-days post culture initiation, and subsequent release until the end of culture (5 days post culture initiation), as well as compared to the corresponding cytokine release by the BCG infected chronic traditional cell culture control (monolayer) and autologous CD3^+^ T cells prior to addition to the AM culture. Each datapoint represents the median of three individual experiments, with BCG infection occurring on day 1 after culture initiation (the day of culture initiation is considered as day 0). The Day 2 inserts for each cytokine depict the differences between cytokines released by uninfected and BCG infected AM 2-days post culture initiation, i.e. 1-day post infection (Each datapoint represents a single individual from the three individual experiments assessed).

### 3D adaptive granuloma structures regulate mycobacterial replication

When total cell numbers from patient samples were not limiting, CFU were measured for each granuloma structure using the cell lysate (Figure 12). For the purposes of this pilot study, we were able to measure the initial uptake by AM after the 4-hour infection from two of the participants which allowed for an indication of bacterial uptake prior to the long-term culture of the 3D granuloma structures (p = 0.2) and traditional cell culture controls (p = 0.8). Due to a lack of adequate cell numbers, the BCG infected innate traditional cell culture control had a single datapoint only and was therefore excluded from analysis. Interestingly, the BCG infected adaptive 3D granuloma displayed a trend for improved bacterial control compared to the BCG infected traditional cell culture control, however with few datapoints available, this was not significant (p = 0.7). It would also appear that for some individuals, the 3D spheroid granulomas are more protective whereas others show the same level of containment between traditional cultures and 3D spheroid granulomas. Increasing the number of datapoints available for CFU count will be of value for determining the levels of significance between bacterial uptake and level of control, but it shows promise that the adaptive 3D granuloma structure shows a trend for improved bacterial control over the traditional cell culture control conditions.

**Figure 12:**
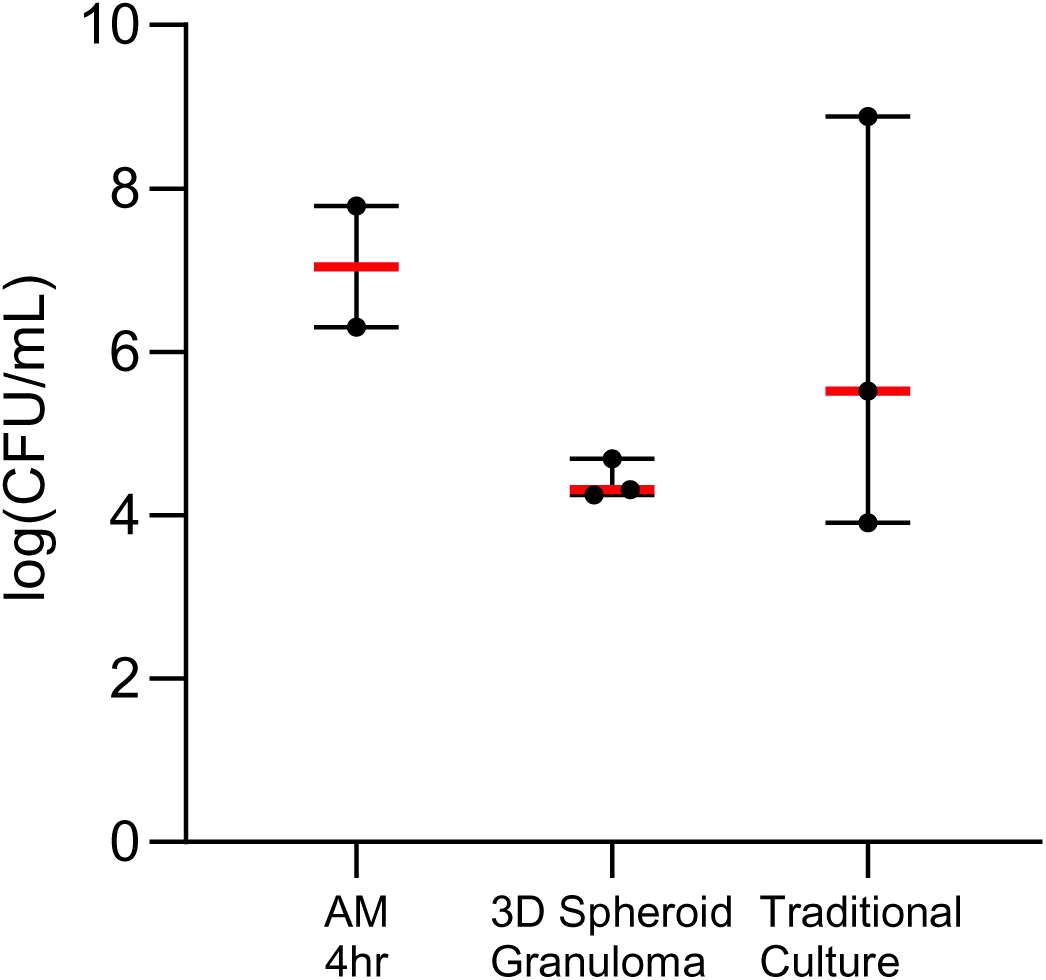
BCG CFU were measured using the cell lysate of 3D spheroid granuloma structures and traditional cell culture control cultures (traditional cultures) and compared to the initial bacterial uptake of BCG into AM. Data were log-transformed prior to plotting.

### Mycobacterial infection and spheroid configuration both alter the gene expression of cells in 3D Spheroid granulomas

We evaluated the ability to extract quality RNA from individually dissociated 3D spheroid granulomas and conducted bulk RNA sequencing on the cells. Total RNA was extracted from eleven available samples, but two did not meet the total RNA requirement for acquisition on the MGISEQ-2000; therefore, only nine samples were used for further evaluations (table in Figure 13). FASTQC was used to assess the quality of the 18 FASTQ files generated, and manual inspection of the quality score graphs of all 18 HTML reports showed that the lowest 10^th^ percentile value for any base at any position was 26; in all cases the software issued a passing grade (Figure 14a, b, d, e). Another important quality metric to consider is the proportion of bases seen at each position. All 18 reads (nine forward and nine reverse reads) showed erratic behaviour in the first 10-13 bases before a transition to a smooth curve for the rest of the read (Figure 14c, f). For RNA-Seq data this is expected and is a result dependent on the specific library kit which was used. As a result, further cleaning, or processing of the raw FASTQ files (for example, trimming of adaptor sequences) was not required.

**Figure 13:**
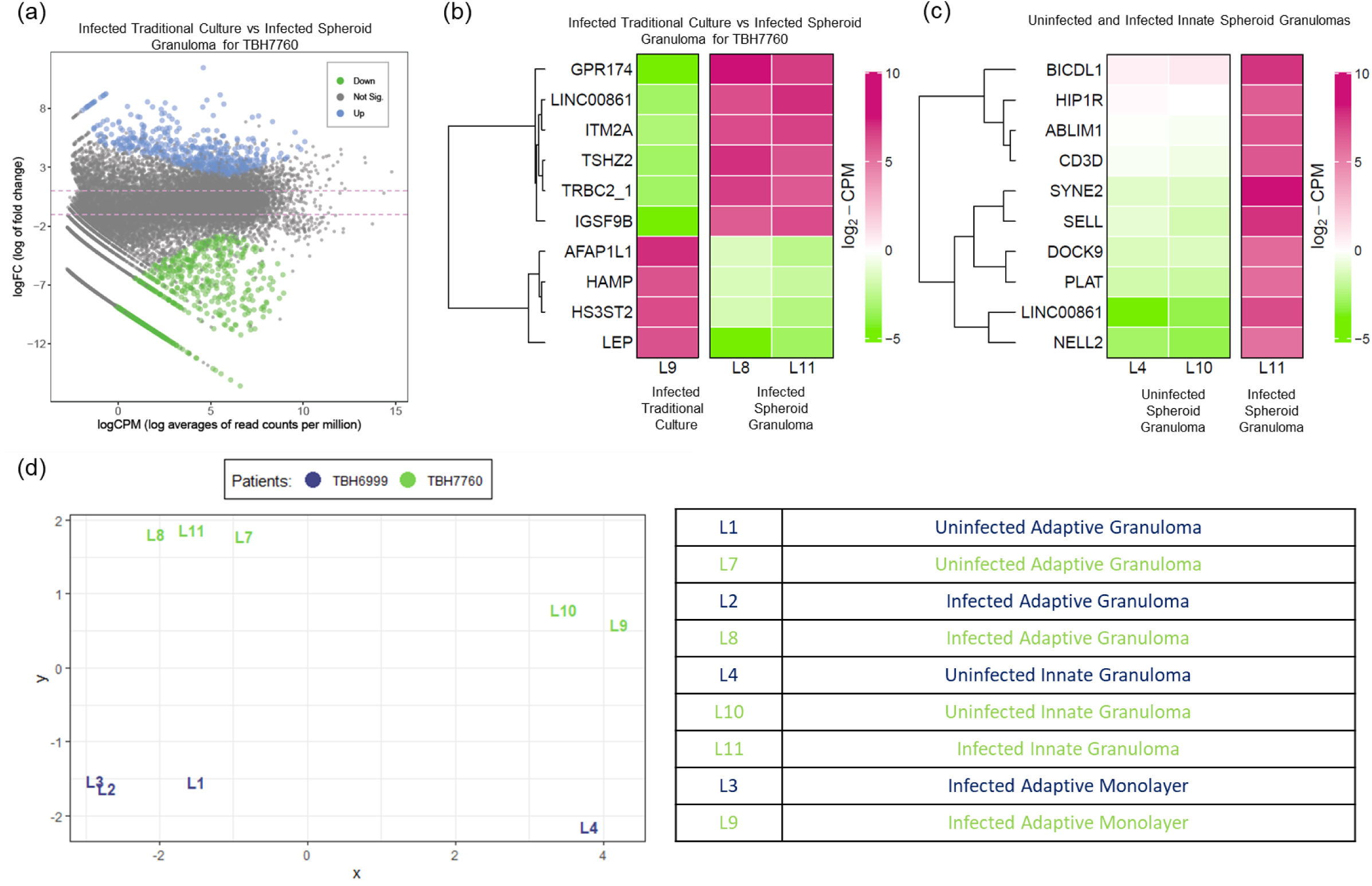
RNA Sequencing relative gene expression results displaying (a) a Mean-Difference (MD) plot of the differential gene expression between BCG Infected traditional control cultures and 3D spheroid granulomas (The distance between any two points is the leading log-fold change between those samples. The leading log-fold change is the root mean square average of the largest log-2 fold-change between those samples.), (b) a heatmap of the differential gene expression of BCG infected cells in traditional culture (labelled “Control”) vs corresponding infected cells in 3D spheroid granulomas, (c) a heatmap of the differential gene expression in BCG infected innate 3D spheroid granulomas vs uninfected innate 3D spheroid granulomas, and (d) a multi-dimensional scaling plot of all datapoints from a smoker (green) vs non-smoker (blue). Descriptions of each datapoint are given in the bottom right-hand table.

**Figure 14:**
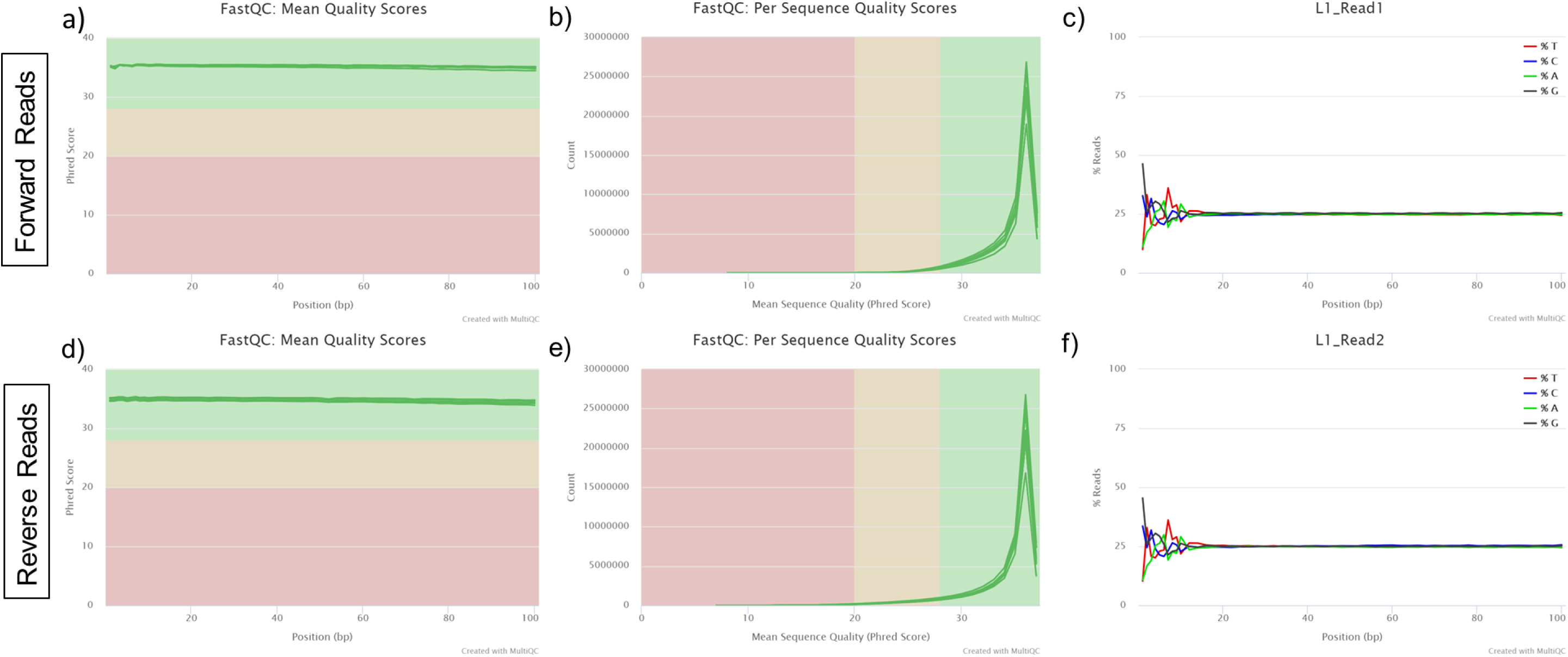
Representative FASTQC quality control results of the nine forward (a-c) and nine reverse (d-f) reads for the nine patient samples run on the MGISEQ-2000. Representative results include (a) the average quality score per base for all nine forward reads, (b) forward read quality score per sequence, (c) an example of the proportion of bases seen at each position for the forward reads where the first 10-13 bases show erratic behaviour, (d) the average quality score per base for all nine reverse reads, (e) reverse read quality score per sequence, and (f) an example of the proportion of bases seen at each position for the reverse reads where the first 10-13 bases show erratic behaviour.

The geometrical impact of 3D granulomas on the biological responses of immune cells, remains unexplored. We therefore compared the gene expression profile of the 3D spheroid granuloma to the corresponding human subject’s cells (matched cell origin, types, ratios, numbers) in traditional cell culture. Results from this demonstrate that cells in 3D spheroid granulomas up-regulate 640 genes and down-regulate 523 genes, when compared to those in traditional culture (Figure 13a). Loose inferences based on the top ten differentially expressed genes (Figure 13b) suggest that 3D spheroid granulomas could augment the transcription of proteins related to inhibitory synapse development, a neuropeptide receptor, a zinc finger transcription factor, and T cell activation, but curb those related to leptin, hepcidin, actin filament proteins and a heparan sulphate-glucosamine enzyme. In total, we identified 138 genes with significant differential expression between BCG-infected 3D spheroid granulomas and BCG-infected traditional culture. These 138 differentially regulated transcripts were functionally annotated to gain an overview of the biological pathway regulation using GO enrichment analysis. The REVIGO resource was used to summarise and visualize the most enriched GO terms for biological processes (Figure 15A), molecular function (Figure 15B) and cellular components (Figure 15C) in an interactive graph. Differential gene expression was also observed between uninfected and BCG-infected 3D spheroid granulomas when innate 3D spheroid granulomas were investigated (Figure 13c), while a multi-dimensional scaling plot of all datapoints demonstrated differences between both participants from which RNA-Seq samples were available as observed by the grouping in the first component between the two different participants (Figure 13d; smoker (green) vs non-smoker (blue)).

**Figure 15:**
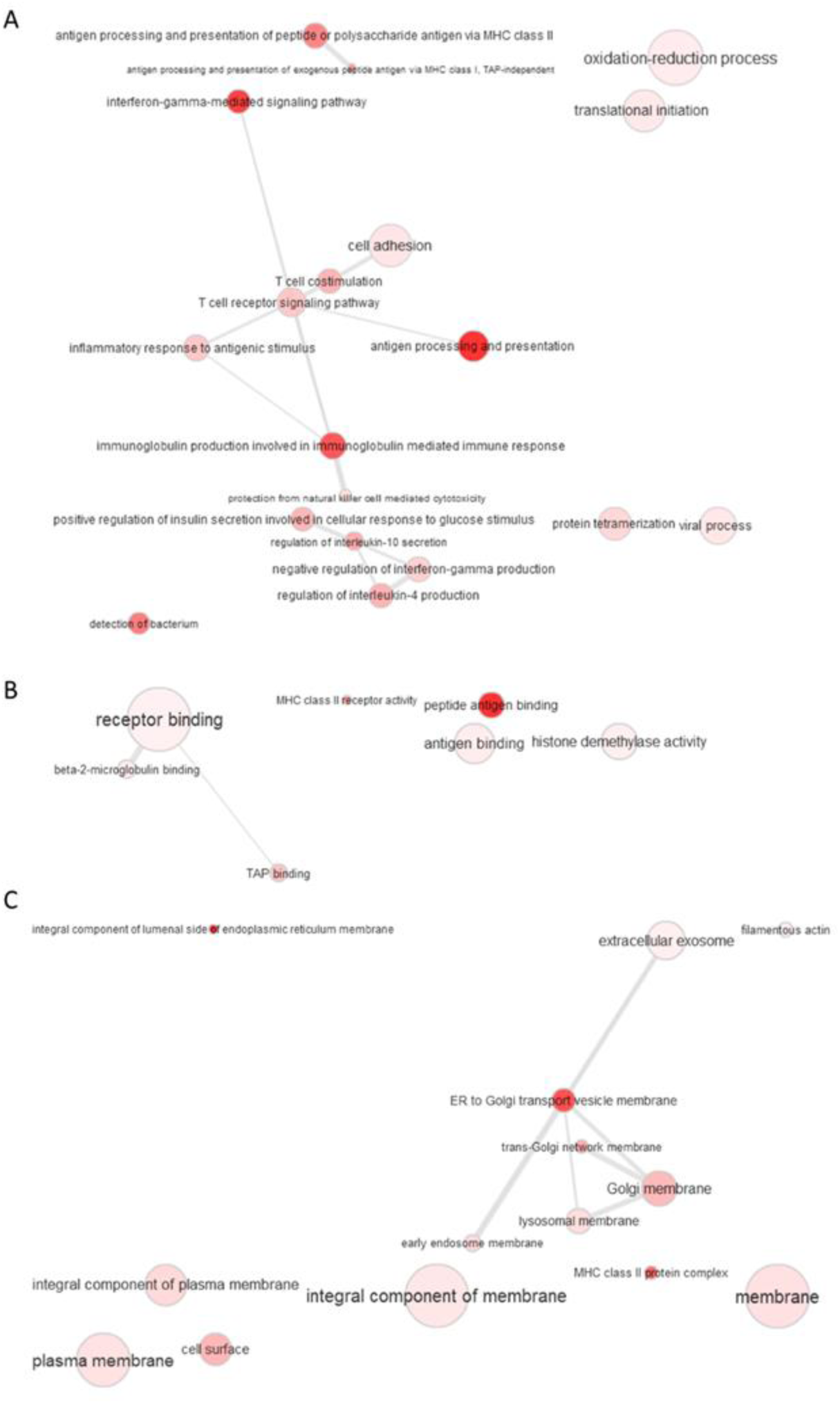
Functional network map of the GO term enrichment analysis for the 138 differentially regulated proteins identified during RNA seq analysis from comparing BCG-infected spheroid granulomas between participants (one a smoker, and one not). GO terms retrieved by the Database for Annotation, Visualization and Integrated Discovery (DAVID) and visualized in ReviGO clustered enriched terms according to (A) biological processes, (B) molecular function and (c) cellular component. Each node corresponds to a single representative GO term for all related sibling and child terms. Highly similar GO terms are linked by edges in the graph where line width indicates the degree of similarity. Bubble colour intensity indicates the p-value and bubble size indicates the frequency of the GO term in the GOA database. Force-directed layout algorithm was used to keep similar nodes together.

**Figure 16:**
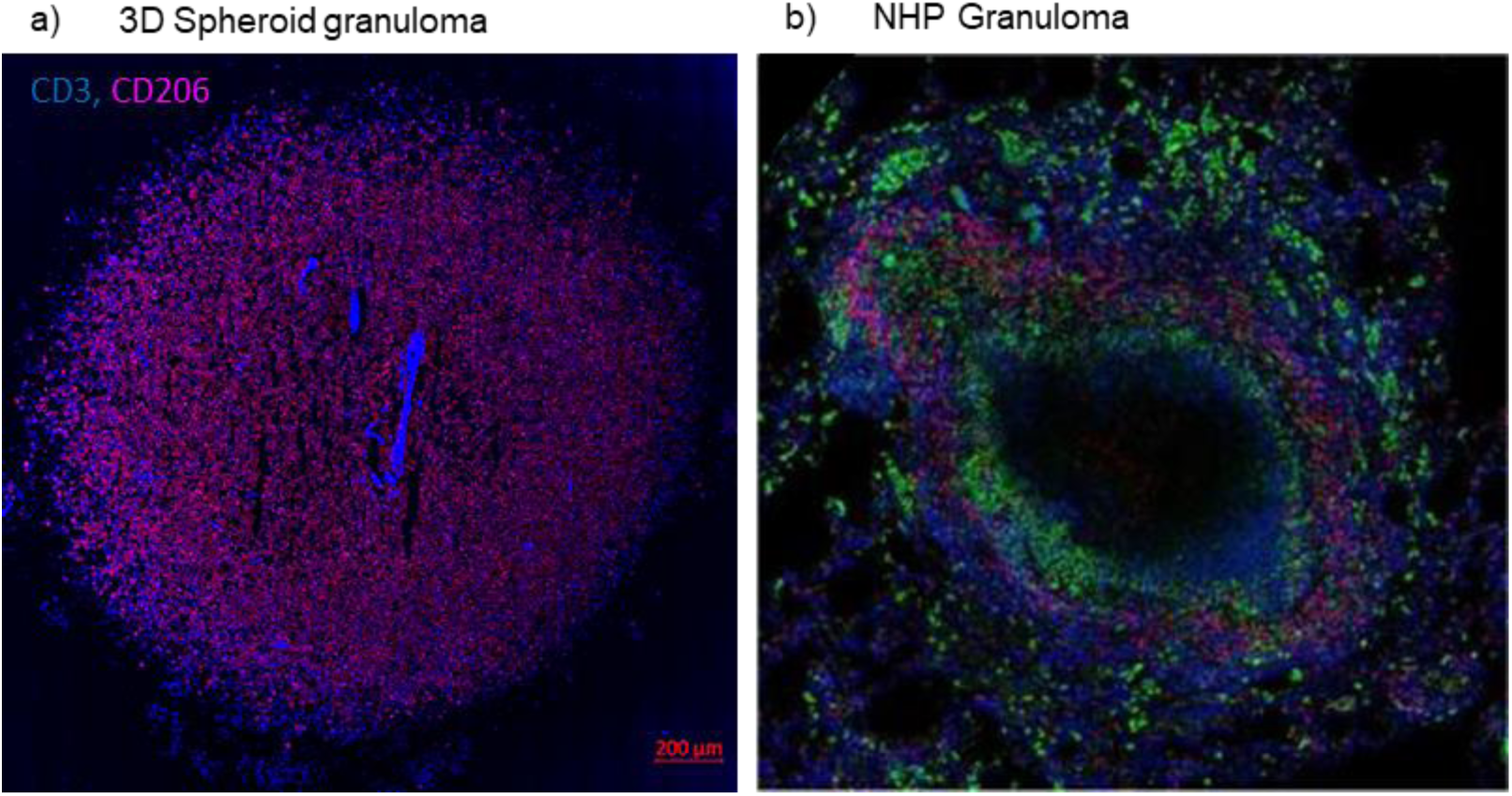
Our 3D in vitro TB granuloma structures show similar structural and cellular composition based on (a) immunofluorescence staining with antibodies for CD3^+^ T-cells (blue, V450) and alveolar macrophages (red, PE-CF594) to (b) published in vivo TB lung granulomas of non-human primates, stained with antibodies for CD3^+^ T-cells (red) and CD68^+^ macrophages (green), surrounding the necrotic center (unstained). Adapted from Flynn *et* al. 2015 with permission (license number: 4967651365284) (Flynn et al., 2015).

While still very preliminary, our RNA-Seq data demonstrates the ability to successfully isolate and sequence high-quality RNA from the 3D spheroid structures after prolonged culture and infection. The preliminary data alludes to differential behaviour of host immune cells based on structural organisation and conformation upon BCG-infection, as observed in the differences between traditional cell culture and the 3D spheroid structures, which may, in future, lead to the possibility of demonstrating that traditional cell culture methods do not accurately reflect responses occurring at a granuloma level during mycobacterial infection. Since this study developed the method of generating 3D spheroid granulomas, future studies will include the comparison of larger numbers of participants to investigate this hypothesis.

## DISCUSSION

Human responses to *M.tb* infection range in complexity, while the heterogeneity of TB disease is unprecedented and challenging to model. Most infected individuals can mount a protective immune response to control infection, resulting in a large proportion of these individuals achieving sterilizing cure. Yet a small proportion will develop active TB disease and some retain a latent infection where they are able to control infection but not achieve sterilizing cure (Barry et al., 2009; Kapoor et al., 2013; O’Garra et al., 2013). Even within individuals, granulomatous lesions present as dynamic and localised microenvironments within the lung, each with their own unique organisation and ability to control infection, ranging from sterilizing cure to immune failure (Barry et al., 2009; Ehlers and Schaible, 2013; Kiran et al., 2016; Lin and Flynn, 2018).

A major knowledge gap is the exact nature, function and spatial organization of immune cells constituting protective granulomas. A hallmark of progression to active TB disease is changes to granuloma physiology, such as an increase in granuloma number and distribution; as well as changes in granuloma function, such as poor *M.tb* replication control and development of central necrosis and cavitation. These signify the host’s inability to eliminate bacilli and are indicative of failed immunity. As an added complexity, differences exist in the rate and trajectory of granuloma progression, which is determined by features such as host immunity and bacterial virulence (Guirado and Schlesinger, 2013; Russell et al., 2009; Silva Miranda et al., 2012). Therefore, TB patients simultaneously harbour a range of granulomas, comprising a spectrum of solid non-necrotizing, necrotic and caseous granulomas, each with its distinct kinetics. Although the overall clinical picture is likely defined by a set of the most poorly performing granulomas in a particular host, granulomas from a single individual vary considerably in cellular composition, cytokine profile, morphology, immune phenotype, and bacterial burden. It is therefore important to describe host responses at the individual granuloma level.

While animal models have contributed significantly to our understanding of the mammalian immune system, numerous examples have shown that laboratory species do not faithfully or in full, mimic human immunity or TB disease (Fonseca et al., 2017; Singh and Gupta, 2018; Zhan et al., 2017). A major deficiency in many mouse models, for example, is the lack of necrotic lung lesions, which is the pathological hallmark advanced granulomas (Kramnik and Beamer, 2016). Other species models such as rabbits and non-human primates, have been useful in studying lung pathologies, but these models are expensive, laborious to maintain and biological reagents are limited. The heterogeneity in host responses to *M.tb* infection is currently more readily investigated in animal models such as cynomolgus macaques, with findings that are translatable to human TB. As such, our understanding of the structure and function of human TB granulomas is inferred from features reported for NHP TB granulomas. Our 3D adaptive spheroid granuloma has here been demonstrated to resemble the structure and features reported for NHP TB granulomas, as is evidenced by the spatial arrangement of an autologous T-cell cuff surrounding an AM core (Figure 15; (Flynn et al., 2015; Kauffman et al., 2018; Wong et al., 2018). This model allows for manipulation of biological targets, and importantly, captures *in vivo* characteristics, such as cellular phenotype and spatial organization as observed in *in vivo* human and NHP granulomas. This could have multiple benefits over traditional culture methods and enable assessment of host responses to *M.tb* in the context of intricate cellular interactions and visualization of granuloma organization through 3D and quantitative analysis. The development of granuloma models that accurately reflect the major pathophysiological conditions existing in the spectrum of *in vivo* human pulmonary TB granulomas, both kinetically and in different clinical phases of infection, is imperative. Ideally, this would require the use of cells derived from patients within the spectrum of TB disease and be retrieved from the site-of-disease to fully recapitulate physiological events occurring within the human lung when challenged with *M.tb.* We have established a novel, laboratory-based, biologically relevant platform for generating patient-derived 3D spheroid granulomas mimicking human TB. The platform enables analysis of genomic, epigenetic, immunologic, structural, pathogen and treatment-specific aspects of immune cells during granuloma evolution, to resemble human pulmonary TB lesions more closely. The 3D spheroid granuloma model is assembled as a single, organized structure in a culture well, consisting of human lung-derived AM, surrounded by layers of autologous, peripherally recruited T-cells. This model has the potential to replicate characteristics observed during granuloma evolution (Driver et al., 2012; Lenaerts et al., 2015) such as hypoxia, nutrient concentration gradients, and for in-depth mechanistic analysis of crucial lung granulomatous features observed in the spectrum from latent to active TB.

Granulomas are known to be dynamic, organized structures with gene expression and epigenetic profiles correlating with lesion type and developmental trajectory. Considering the limitations associated with *in vivo* human research, few studies have explored the immune environment within human lung granulomas (Kaplan et al., 2003; Kim et al., 2010; Mattila et al., 2013). Results have however demonstrated that pooled analysis of different human lung granuloma types may underestimate differences in gene expression specific to each lesion, revealing a higher number of significantly differentially expressed genes in fibrotic nodules, compared to the cavitary granulomas (Subbian et al., 2015). Thus, a distinct gene expression pattern was observed for each granuloma type or stage, and increased methylation was found in granulomas of patients with more severe disease (Gautam et al., 2014; Mehra et al., 2013; Yang et al., 2019). Our granuloma model has alluded to these observations, with the extraction of good quality RNA from this model proving that structures generated from individuals along the spectrum of disease are likely capable of providing detailed gene expression profiles for the various stages of infection.

The first guiding 3D model described for *M.tb* infection was described by Puissegur *et al*. in 2004 and has helped direct current model strategies (Puissegur et al., 2004). Since then, a number of models have been established with varied success rates (reviewed by Elkington *et al.,* 2019), and with little focus being placed on primary human cells obtained from the site of disease, one of the proposed requirements of an optimal model (Elkington et al., 2019; Nunes et al., 2019). Most popularly, *in vitro* granuloma-like cell aggregates established using *M.tb* infected PBMC have shown promise in generating cellular aggregates which mimic granuloma formation (Crouser et al., 2017; Guirado et al., 2015). While easy to establish and high throughput, this model of infected PBMC, form multiple structures within a culture well, each structure at a different “stage” of granuloma development, which may limit some aspects of granuloma investigations (Silva Miranda et al., 2012). PBMC granuloma models also do not accurately represent the immune cell milieu and composition, or the defined organizational structure, as observed for the 3D spheroid model. In addition to this, we have demonstrated with various molecular tools the validity of employing such a model for the investigation of *M.tb* infection without the limitations of traditional cell culture methods and the need to acquire entire *in vivo* structures from procedures such as biopsies. The most recently published *in vitro* granuloma model example using biopsy samples includes an adult stem cell-derived airway organoid developed from cells retrieved from human lung biopsies (Sachs et al., 2019). The limitation of such a model is that it requires the need for difficult to obtain sample types, and complex organoid generation methods which do not include the use of primary phagocytes such as alveolar macrophages which are essential for the initial control of *M.tb* infection. Another important consideration for our model is that granuloma formation is not limited to *M.tb* infection, but also occurs in several chronic infections including Schistosoma spp., *S. enterica* and *L. monocytogenes*, and non-infectious diseases like sarcoidosis (Crouser et al., 2017), which together cause millions of deaths. Our granuloma model could therefore be adapted to depict these disease-specific granulomatous features.

Our model is not without its limitations, however. The autofluorescent nature of alveolar macrophages from the lungs of individuals in certain regions like Cape Town, South Africa, display high carbon particulate matter which results in autofluorescence. We have demonstrated that this limitation can be circumvented; alternatively, molecular techniques like CyTOF could be used to investigate both the AM and autologous CD3^+^ T cell fractions together, without the need to consider autofluorescence, as has previously been demonstrated by our group (Young et al., 2019). The use of the magnetic levitation drive, for one, prevents the unrestricted movement of NanoShuttle^TM^-labelled AM until the core is stable, and therefore features of early AM movement could be missed. This model was established using BCG as a model for TB owing to the benefits of being able to work with this organism outside of a BSL3 environment. Considering the physiological differences between BCG and H37Rv, the virulent laboratory strain of *M.tb*, this model needs to be validated in a setting whereby H37Rv is used as the infectious agent instead of BCG. We are confident, however, that the model we have established using BCG as a model organism for *M.tb* will be translatable for not only *M.tb*, but other pulmonary pathogens which result in the formation of pulmonary granulomas. As such, the results generated in this study during the establishment of the 3D spheroid *in vitro* granuloma model should function to inform the scientific community of the possibilities for which this model can be adapted.

Finally, our 3D spheroid *in vitro* granuloma model still requires comparative assessments to *in vivo* granulomas with respect to the cellular components, cell phenotype, molecular and epigenetic interactions, patterns of cytokine and chemokine secretion, mycobacterial dormancy, and subcellular-dissemination and ultimately and impact on clinical outcome. These are currently ongoing through planned human- and non-human primate studies. Additionally, we are currently conducting a study whereby we have begun adding additional cell types, such as B cells and myeloid-derived suppressor cells (MDSCs), to the 3D spheroid granuloma as one would expect to see in a developing granuloma (Agrawal et al., 2018; Flynn et al., 2011; Obregón-Henao et al., 2013). With that said, the current model would provide insights into host-mycobacterial interactions at stages too early to address within such *in vivo* models and eventually, serve as preferred platform for initial pre-clinical testing of TB vaccine and drug candidates.

## ACKNOWLEDGEMENTS

The authors acknowledge the financial support from the European & Developing Countries Clinical Trials Partnership (EDCTP; CDF1546) and International Collaborations in Infectious Disease Research (ICIDR): Biology and Biosignatures of anti-TB Treatment Response (5U01IA115619/03). The authors would also like to acknowledge the doctors and nursing staff of Ward A5, Pulmonology Division of Tygerberg Academic Hospital, especially Sr Lauren Benting, for their willingness to assist in the collection of these samples for research purposes. A special thanks also goes to Sr Ruth Wilson, a research nurse from the CLIME laboratory in the Department of Molecular Biology and Human Genetics, Stellenbosch University, for recruiting the participants for this study, without whom this study would not have been possible. Lastly, the authors would like to thank Tracey Jooste from the South African Medical Research Council (SAMRC) Genomics Centre for her expertise in RNA-Sequencing.

## AUTHOR CONTRIBUTIONS STATEMENT

LAK performed all the experiments necessary for the development and construction of the 3D *in vitro* spheroid granulomas, performed all the downstream processing of the structures (excluding the RNA Sequencing and immunofluorescence staining) and wrote the manuscript. CGGB performed all confocal microscopy experiments (including the optimisation) for the visualisation of the 3D *in vitro* spheroid granulomas, with invaluable assistance from DL and SC during the troubleshooting and optimisation of the immunofluorescence staining of the structures. CGGB also assisted with the writing and editing of the manuscript and performed the pathway analysis for the RNA sequencing data. NDP conceptualised the 3D *in vitro* granuloma model, designing the experiments with LAK, and assisted with the writing, review and editing of the manuscript. CK kindly performed the RNA Sequencing, while BG and MM analysed the FASTQ data, including performing all the quality control checks necessary for proceeding with the analysis. GW reviewed the manuscript and was part of the initial conceptualisation of the project.

## REFERENCES

1. Agrawal, N., Streata, I., Pei, G., Weiner, J., Kotze, L., Bandermann, S., Lozza, L., Walzl, G., du Plessis, N., Ioana, M., Kaufmann, S.H.E., Dorhoi, A., 2018. Human Monocytic Suppressive Cells Promote Replication of Mycobacterium tuberculosis and Alter Stability of in vitro Generated Granulomas. Front. Immunol. 9, 2417. https://doi.org/10.3389/fimmu.2018.02417

2. Barry, C.E., Boshoff, H.I., Dartois, V., Dick, T., Ehrt, S., Flynn, J., Schnappinger, D., Wilkinson, R.J., Young, D., 2009. The spectrum of latent tuberculosis: rethinking the biology and intervention strategies. Nat. Rev. Microbiol. 7, 845–855. https://doi.org/10.1038/nrmicro2236

3. Belton, M., Brilha, S., Manavaki, R., Mauri, F., Nijran, K., Hong, Y.T., Patel, N.H., Dembek, M., Tezera, L., Green, J., Moores, R., Aigbirhio, F., Al-Nahhas, A., Fryer, T.D., Elkington, P.T., Friedland, J.S., 2016. Hypoxia and tissue destruction in pulmonary TB. Thorax 71, 1145–1153. https://doi.org/10.1136/thoraxjnl-2015-207402

4. Canetti, G., 1955. The Tubercle Bacillus in the Pulmonary Lesion of Man: Histobacteriology and Its Bearing on the Therapy of Pulmonary Tuberculosis. Springer Publishing Company.

5. Cohen, S.B., Gern, B.H., Delahaye, J.L., Adams, K.N., Plumlee, C.R., Winkler, J.K., Sherman, D.R., Gerner, M.Y., Urdahl, K.B., 2018. Alveolar Macrophages Provide an Early Mycobacterium tuberculosis Niche and Initiate Dissemination. Cell Host Microbe 24, 439–446.e4. https://doi.org/10.1016/j.chom.2018.08.001

6. Crouser, E.D., White, P., Caceres, E.G., Julian, M.W., Papp, A.C., Locke, L.W., Sadee, W., Schlesinger, L.S., 2017. A Novel In Vitro Human Granuloma Model of Sarcoidosis and Latent Tuberculosis Infection. Am. J. Respir. Cell Mol. Biol. 57, 487–498. https://doi.org/10.1165/rcmb.2016-0321OC

7. Davis, J.M., Ramakrishnan, L., 2009. The role of the granuloma in expansion and dissemination of early tuberculous infection. Cell 136, 37–49. https://doi.org/10.1016/j.cell.2008.11.014

8. Driver, E.R., Ryan, G.J., Hoff, D.R., Irwin, S.M., Basaraba, R.J., Kramnik, I., Lenaerts, A.J., 2012. Evaluation of a Mouse Model of Necrotic Granuloma Formation Using C3HeB/FeJ Mice for Testing of Drugs against Mycobacterium tuberculosis. Antimicrob. Agents Chemother. 56, 3181–3195. https://doi.org/10.1128/AAC.00217-12

9. Duval, K., Grover, H., Han, L.-H., Mou, Y., Pegoraro, A.F., Fredberg, J., Chen, Z., 2017. Modeling Physiological Events in 2D vs. 3D Cell Culture. Physiology 32, 266–277. https://doi.org/10.1152/physiol.00036.2016

10. Ehlers, S., Schaible, U.E., 2013. The Granuloma in Tuberculosis: Dynamics of a Host– Pathogen Collusion. Front. Immunol. 3 . https://doi.org/10.3389/fimmu.2012.00411

11. Elkington, P., Lerm, M., Kapoor, N., Mahon, R., Pienaar, E., Huh, D., Kaushal, D., Schlesinger, L.S., 2019. In Vitro Granuloma Models of Tuberculosis: Potential and Challenges. J. Infect. Dis. 219, 1858–1866. https://doi.org/10.1093/infdis/jiz020

12. Flynn, J.L., Chan, J., Lin, P.L., 2011. Macrophages and control of granulomatous inflammation in tuberculosis. Mucosal Immunol. 4, 271–278. https://doi.org/10.1038/mi.2011.14

13. Flynn, J.L., Gideon, H.P., Mattila, J.T., Lin, P.L., 2015. Immunology studies in non-human primate models of tuberculosis. Immunol. Rev. 264, 60–73. https://doi.org/10.1111/imr.12258

14. Fonseca, K.L., Rodrigues, P.N.S., Olsson, I.A.S., Saraiva, M., 2017. Experimental study of tuberculosis: From animal models to complex cell systems and organoids. PLOS Pathog. 13, e1006421. https://doi.org/10.1371/journal.ppat.1006421

15. Gautam, U.S., Mehra, S., Ahsan, M.H., Alvarez, X., Niu, T., Kaushal, D., 2014. Role of TNF in the altered interaction of dormant Mycobacterium tuberculosis with host macrophages. PloS One 9, e95220. https://doi.org/10.1371/journal.pone.0095220

16. Gil, O., Díaz, I., Vilaplana, C., Tapia, G., Díaz, J., Fort, M., Cáceres, N., Pinto, S., Caylà, J., Corner, L., Domingo, M., Cardona, P.-J., 2010. Granuloma encapsulation is a key factor for containing tuberculosis infection in minipigs. PloS One 5, e10030. https://doi.org/10.1371/journal.pone.0010030

17. Guirado, E., Mbawuike, U., Keiser, T.L., Arcos, J., Azad, A.K., Wang, S.-H., Schlesinger, L.S., 2015. Characterization of Host and Microbial Determinants in Individuals with Latent Tuberculosis Infection Using a Human Granuloma Model. mBio 6. https://doi.org/10.1128/mBio.02537-14

18. Guirado, E., Schlesinger, L.S., 2013. Modeling the Mycobacterium tuberculosis Granuloma - the Critical Battlefield in Host Immunity and Disease. Front. Immunol. 4, 98. https://doi.org/10.3389/fimmu.2013.00098

19. Haisler, W.L., Timm, D.M., Gage, J.A., Tseng, H., Killian, T.C., Souza, G.R., 2013. Three-dimensional cell culturing by magnetic levitation. Nat. Protoc. 8, 1940–1949. https://doi.org/10.1038/nprot.2013.125

20. Herrmann, K., Pistollato, F., Stephens, M., 2019. Beyond the 3Rs: Expanding the Use of Human-Relevant Replacement Methods in Biomedical Research. Biomed. Res. Altern. Methods Collect.

21. Hirschhaeuser, F., Menne, H., Dittfeld, C., West, J., Mueller-Klieser, W., Kunz-Schughart, L.A., 2010. Multicellular tumor spheroids: an underestimated tool is catching up again. J. Biotechnol. 148, 3–15. https://doi.org/10.1016/j.jbiotec.2010.01.012

22. Huang, D.W., Sherman, B.T., Lempicki, R.A., 2009a. Systematic and integrative analysis of large gene lists using DAVID bioinformatics resources. Nat. Protoc. 4, 44–57. https://doi.org/10.1038/nprot.2008.211

23. Huang, D.W., Sherman, B.T., Lempicki, R.A., 2009b. Bioinformatics enrichment tools: paths toward the comprehensive functional analysis of large gene lists. Nucleic Acids Res. 37, 1–13. https://doi.org/10.1093/nar/gkn923

24. Kaplan, G., Post, F.A., Moreira, A.L., Wainwright, H., Kreiswirth, B.N., Tanverdi, M., Mathema, B., Ramaswamy, S.V., Walther, G., Steyn, L.M., Barry, C.E., Bekker, L.-G., 2003. Mycobacterium tuberculosis growth at the cavity surface: a microenvironment with failed immunity. Infect. Immun. 71, 7099–7108. https://doi.org/10.1128/iai.71.12.7099-7108.2003

25. Kapoor, N., Pawar, S., Sirakova, T.D., Deb, C., Warren, W.L., Kolattukudy, P.E., 2013. Human granuloma in vitro model, for TB dormancy and resuscitation. PloS One 8, e53657. https://doi.org/10.1371/journal.pone.0053657

26. Kauffman, K.D., Sallin, M.A., Sakai, S., Kamenyeva, O., Kabat, J., Weiner, D., Sutphin, M., Schimel, D., Via, L., Barry, C.E., Wilder-Kofie, T., Moore, I., Moore, R., Barber, D.L., 2018. Defective positioning in granulomas but not lung-homing limits CD4 T-cell interactions with Mycobacterium tuberculosis-infected macrophages in rhesus macaques. Mucosal Immunol. 11, 462–473. https://doi.org/10.1038/mi.2017.60

27. Kim, M.-J., Wainwright, H.C., Locketz, M., Bekker, L.-G., Walther, G.B., Dittrich, C., Visser, A., Wang, W., Hsu, F.-F., Wiehart, U., Tsenova, L., Kaplan, G., Russell, D.G., 2010. Caseation of human tuberculosis granulomas correlates with elevated host lipid metabolism. EMBO Mol. Med. 2, 258–274. https://doi.org/10.1002/emmm.201000079

28. Kiran, D., Podell, B.K., Chambers, M., Basaraba, R.J., 2016. Host-directed therapy targeting the Mycobacterium tuberculosis granuloma: a review. Semin. Immunopathol. 38, 167–183. https://doi.org/10.1007/s00281-015-0537-x

29. Kramnik, I., Beamer, G., 2016. Mouse models of human TB pathology: roles in the analysis of necrosis and the development of host-directed therapies. Semin. Immunopathol. 38, 221–237. https://doi.org/10.1007/s00281-015-0538-9

30. Lenaerts, A., Barry, C.E., Dartois, V., 2015. Heterogeneity in tuberculosis pathology, microenvironments and therapeutic responses. Immunol. Rev. 264, 288–307. https://doi.org/10.1111/imr.12252

31. Lillie, R.D., 1965. Histopathologic technic and practical histochemistry., 3rd ed. ed. Blakiston Division, McGraw-Hill, New York.

32. Lin, P.L., Flynn, J.L., 2018. The End of the Binary Era: Revisiting the Spectrum of Tuberculosis. J. Immunol. 201, 2541–2548. https://doi.org/10.4049/jimmunol.1800993

33. Lin, P.L., Pawar, S., Myers, A., Pegu, A., Fuhrman, C., Reinhart, T.A., Capuano, S.V., Klein, E., Flynn, J.L., 2006. Early events in Mycobacterium tuberculosis infection in cynomolgus macaques. Infect. Immun. 74, 3790–3803. https://doi.org/10.1128/IAI.00064-06

34. Mattila, J.T., Ojo, O.O., Kepka-Lenhart, D., Marino, S., Kim, J.H., Eum, S.Y., Via, L.E., Barry, C.E., Klein, E., Kirschner, D.E., Morris, S.M., Lin, P.L., Flynn, J.L., 2013. Microenvironments in tuberculous granulomas are delineated by distinct populations of macrophage subsets and expression of nitric oxide synthase and arginase isoforms. J. Immunol. Baltim. Md 1950 191, 773–784. https://doi.org/10.4049/jimmunol.1300113

35. Mehra, S., Alvarez, X., Didier, P.J., Doyle, L.A., Blanchard, J.L., Lackner, A.A., Kaushal, D., 2013. Granuloma Correlates of Protection Against Tuberculosis and Mechanisms of Immune Modulation by Mycobacterium tuberculosis. J. Infect. Dis. 207, 1115–1127. https://doi.org/10.1093/infdis/jis778

36. Nunes, A.S., Barros, A.S., Costa, E.C., Moreira, A.F., Correia, I.J., 2019. 3D tumor spheroids as in vitro models to mimic in vivo human solid tumors resistance to therapeutic drugs. Biotechnol. Bioeng. 116, 206–226. https://doi.org/10.1002/bit.26845

37. Obregón-Henao, A., Henao-Tamayo, M., Orme, I.M., Ordway, D.J., 2013. Gr1intCD11b+ Myeloid-Derived Suppressor Cells in Mycobacterium tuberculosis Infection. PLoS ONE 8. https://doi.org/10.1371/journal.pone.0080669

38. O’Garra, A., Redford, P.S., McNab, F.W., Bloom, C.I., Wilkinson, R.J., Berry, M.P.R., 2013. The Immune Response in Tuberculosis. Annu. Rev. Immunol. 31, 475–527. https://doi.org/10.1146/annurev-immunol-032712-095939

39. Pagán, A.J., Ramakrishnan, L., 2014. Immunity and Immunopathology in the Tuberculous Granuloma. Cold Spring Harb. Perspect. Med. 5. https://doi.org/10.1101/cshperspect.a018499

40. Parasa, V.R., Rahman, M.J., Hoang, A.T.N., Svensson, M., Brighenti, S., Lerm, M., 2014. Modeling Mycobacterium tuberculosis early granuloma formation in experimental human lung tissue. Dis. Model. Mech. 7, 281–288. https://doi.org/10.1242/dmm.013854

41. Peyron, P., Vaubourgeix, J., Poquet, Y., Levillain, F., Botanch, C., Bardou, F., Daffé, M., Emile, J.-F., Marchou, B., Cardona, P.-J., de Chastellier, C., Altare, F., 2008. Foamy macrophages from tuberculous patients’ granulomas constitute a nutrient-rich reservoir for M. tuberculosis persistence. PLoS Pathog. 4, e1000204. https://doi.org/10.1371/journal.ppat.1000204

42. Puissegur, M.-P., Botanch, C., Duteyrat, J.-L., Delsol, G., Caratero, C., Altare, F., 2004. An in vitro dual model of mycobacterial granulomas to investigate the molecular interactions between mycobacteria and human host cells. Cell. Microbiol. 6, 423– 433. https://doi.org/10.1111/j.1462-5822.2004.00371.x

43. Russell, D.G., Cardona, P.-J., Kim, M.-J., Allain, S., Altare, F., 2009. Foamy macrophages and the progression of the human TB granuloma. Nat. Immunol. 10, 943–948. https://doi.org/10.1038/ni.1781

44. Sachs, N., Papaspyropoulos, A., Zomer-van Ommen, D.D., Heo, I., Böttinger, L., Klay, D., Weeber, F., Huelsz-Prince, G., Iakobachvili, N., Amatngalim, G.D., de Ligt, J., van Hoeck, A., Proost, N., Viveen, M.C., Lyubimova, A., Teeven, L., Derakhshan, S., Korving, J., Begthel, H., Dekkers, J.F., Kumawat, K., Ramos, E., van Oosterhout, M.F., Offerhaus, G.J., Wiener, D.J., Olimpio, E.P., Dijkstra, K.K., Smit, E.F., van der Linden, M., Jaksani, S., van de Ven, M., Jonkers, J., Rios, A.C., Voest, E.E., van Moorsel, C.H., van der Ent, C.K., Cuppen, E., van Oudenaarden, A., Coenjaerts, F.E., Meyaard, L., Bont, L.J., Peters, P.J., Tans, S.J., van Zon, J.S., Boj, S.F., Vries, R.G., Beekman, J.M., Clevers, H., 2019. Long-term expanding human airway organoids for disease modeling. EMBO J. 38, e100300. https://doi.org/10.15252/embj.2018100300

45. Salceda, S., Caro, J., 1997. Hypoxia-inducible factor 1alpha (HIF-1alpha) protein is rapidly degraded by the ubiquitin-proteasome system under normoxic conditions. Its stabilization by hypoxia depends on redox-induced changes. J. Biol. Chem. 272, 22642–22647. https://doi.org/10.1074/jbc.272.36.22642

46. Silva Miranda, M., Breiman, A., Allain, S., Deknuydt, F., Altare, F., 2012. The tuberculous granuloma: an unsuccessful host defence mechanism providing a safety shelter for the bacteria? Clin. Dev. Immunol. 2012, 139127. https://doi.org/10.1155/2012/139127

47. Singh, A.K., Gupta, U.D., 2018. Animal models of tuberculosis: Lesson learnt. Indian J. Med. Res. 147, 456–463. https://doi.org/10.4103/ijmr.IJMR_554_18

48. Souza, G.R., Molina, J.R., Raphael, R.M., Ozawa, M.G., Stark, D.J., Levin, C.S., Bronk, L.F., Ananta, J.S., Mandelin, J., Georgescu, M.-M., Bankson, J.A., Gelovani, J.G., Killian, T.C., Arap, W., Pasqualini, R., 2010. Three-dimensional Tissue Culture Based on Magnetic Cell Levitation. Nat. Nanotechnol. 5, 291–296. https://doi.org/10.1038/nnano.2010.23

49. Subbian, S., Tsenova, L., Kim, M.-J., Wainwright, H.C., Visser, A., Bandyopadhyay, N., Bader, J.S., Karakousis, P.C., Murrmann, G.B., Bekker, L.-G., Russell, D.G., Kaplan, G., 2015. Lesion-Specific Immune Response in Granulomas of Patients with Pulmonary Tuberculosis: A Pilot Study. PLoS ONE 10. https://doi.org/10.1371/journal.pone.0132249

50. Supek, F., Bošnjak, M., Škunca, N., Šmuc, T., 2011. REVIGO Summarizes and Visualizes Long Lists of Gene Ontology Terms. PLOS ONE 6, e21800. https://doi.org/10.1371/journal.pone.0021800

51. Tseng, H., Gage, J.A., Raphael, R.M., Moore, R.H., Killian, T.C., Grande-Allen, K.J., Souza, G.R., 2013. Assembly of a three-dimensional multitype bronchiole coculture model using magnetic levitation. Tissue Eng. Part C Methods 19, 665–675. https://doi.org/10.1089/ten.TEC.2012.0157

52. Tseng, H., Gage, J.A., Shen, T., Haisler, W.L., Neeley, S.K., Shiao, S., Chen, J., Desai, P.K., Liao, A., Hebel, C., Raphael, R.M., Becker, J.L., Souza, G.R., 2015. A spheroid toxicity assay using magnetic 3D bioprinting and real-time mobile device-based imaging. Sci. Rep. 5. https://doi.org/10.1038/srep13987

53. Wolf, A.J., Linas, B., Trevejo-Nuñez, G.J., Kincaid, E., Tamura, T., Takatsu, K., Ernst, J.D., 2007. Mycobacterium tuberculosis infects dendritic cells with high frequency and impairs their function in vivo. J. Immunol. Baltim. Md 1950 179, 2509–2519. https://doi.org/10.4049/jimmunol.179.4.2509

54. Wong, E.A., Joslyn, L., Grant, N.L., Klein, E., Lin, P.L., Kirschner, D.E., Flynn, J.L., 2018. Low Levels of T Cell Exhaustion in Tuberculous Lung Granulomas. Infect. Immun. 86. https://doi.org/10.1128/IAI.00426-18

55. Yang, I.V., Konigsberg, I., MacPhail, K., Li, L., Davidson, E.J., Mroz, P.M., Hamzeh, N., Gillespie, M., Silveira, L.J., Fingerlin, T.E., Maier, L.A., 2019. DNA Methylation Changes in Lung Immune Cells Are Associated with Granulomatous Lung Disease. Am. J. Respir. Cell Mol. Biol. 60, 96–105. https://doi.org/10.1165/rcmb.2018-0177OC

56. Young, C., Ahlers, P., Hiemstra, A.M., Loxton, A.G., Gutschmidt, A., Malherbe, S.T., Walzl, G., Du Plessis, N., Koegelenberg, C.F.N., Kleynhans, L., Ronacher, K., Shaw, J.A., Simon, D., McAnda, S., Swartz, K.C., the SU-IRG consortium, 2019. Performance and immune characteristics of bronchoalveolar lavage by research bronchoscopy in pulmonary tuberculosis and other lung diseases in the Western Cape, South Africa. Transl. Med. Commun. 4, 7. https://doi.org/10.1186/s41231-019-0039-2

57. Zhan, L., Tang, J., Sun, M., Qin, C., 2017. Animal Models for Tuberculosis in Translational and Precision Medicine. Front. Microbiol. 8. https://doi.org/10.3389/fmicb.2017.00717

